# High-Throughput Discovery and Characterization of Viral Transcriptional Effectors in Human Cells

**DOI:** 10.1101/2022.12.16.520835

**Authors:** Connor H. Ludwig, Abby R. Thurm, David W. Morgens, Kevin J. Yang, Josh Tycko, Michael C. Bassik, Britt A. Glaunsinger, Lacramioara Bintu

## Abstract

Viruses encode transcriptional regulatory proteins critical for controlling viral and host gene expression. Given their multifunctional nature and high sequence divergence, it is unclear which viral proteins can affect transcription and which specific sequences contribute to this function. Using a high-throughput assay, we measured the transcriptional regulatory potential of over 60,000 protein tiles across ∼1,500 proteins from 11 coronaviruses and all nine human herpesviruses. We discovered hundreds of new transcriptional effector domains, including a conserved repression domain in all coronavirus Spike homologs, dual activation-repression domains in VIRFs, and an activation domain in six herpesvirus homologs of the single-stranded DNA-binding protein that we show is important for viral replication and late gene expression in KSHV. For the effector domains we identified, we investigated their mechanisms via high-throughput sequence and chemical perturbations, pinpointing sequence motifs essential for function. This work massively expands viral protein annotations, serving as a springboard for studying their biological and health implications and providing new candidates for compact gene regulation tools.

## Introduction

There are more than 200 viruses that infect humans, many of which are known etiological agents of disease^1^ and have been responsible for major global health crises, including the most recent COVID-19 pandemic. Key to this pathogenicity are interactions between viral factors and host cellular machinery^2^. Viruses encode transcriptional regulatory proteins, which are critical for the precise temporal control of viral gene expression and the extensive rewiring of host gene expression programs necessary for creating a cellular environment conducive to productive infection^3^. Viral transcriptional regulators (vTRs) are thus attractive targets for therapeutic intervention^4^.

Given the multifunctional nature of many viral proteins, which have evolved so due to virion and genome size constraints^5^, and their relatively high sequence divergence^6^, it is not clear which viral proteins can affect host transcription. A recently published meta-analysis compiled a census of viral proteins with evidence for nucleic acid binding and/or transcriptional regulation and examined their properties, secondary functions, and genomic targets for the small subset of proteins for which data was available^7^. While this represents the best compilation of vTRs to date, many of the entries within the vTR census lack direct experimental evidence of transcriptional regulation, most of their effector domains have not yet been defined, and the census as a whole likely only represents a fraction of all vTRs due to historical technical limitations that have precluded systematic experimental investigation of transcriptional effector function.

In this study, we use a recently developed high-throughput approach^8^ to test tens of thousands of protein sequences for their effect on gene expression when recruited at reporter genes. This method allows us to identify and characterize viral transcriptional regulators and their effector domains. We start with entries from the vTR census to demonstrate feasibility. We then extend this approach to discover previously undescribed effector domains within the proteins of 11 coronaviruses, including SARS-CoV-2, and all nine human herpesviruses. For the hundreds of effector proteins that we identify, we investigate the sequence determinants of transcriptional regulation, their mechanisms of action using high-throughput measurements, and for a small subset of them their consequences on host gene expression.

### High-throughput identification of activation and repression domains across a curated library of putative viral transcriptional regulators

We have recently developed a high-throughput method (HT-recruit) that allows us to measure the activity of thousands of transcriptional activators and repressors at reporter genes (**Fig. 1A**)^8^. We do this by cloning a library of putative regulators as fusions to the doxycycline (dox)-inducible rTetR DNA binding domain and delivering them to K562 cells by lentivirus at low multiplicity of infection such that each cell contains a single library member. By adding dox, we can recruit candidate effector domains to a minimal promoter (minCMV) to identify activators or to a constitutive promoter (EF1ɑ) to identify repressors (**Fig. 1A**). The reporter genes encode both a fluorescent protein for visualization as well as a surface marker for rapid and robust magnetic separation based on reporter transcriptional state (ON or OFF). Following magnetic separation, we extract genomic DNA from cells in the ON and OFF populations, prepare libraries for next generation sequencing, and compute quantitative enrichment scores for each library member based on their frequencies in the two populations. This method allows us to measure the activity of tens of thousands of candidate effector domains, each 80 amino acids (aa) long (the current limit of DNA synthesis for pooled libraries of this size).

**Fig. 1.**
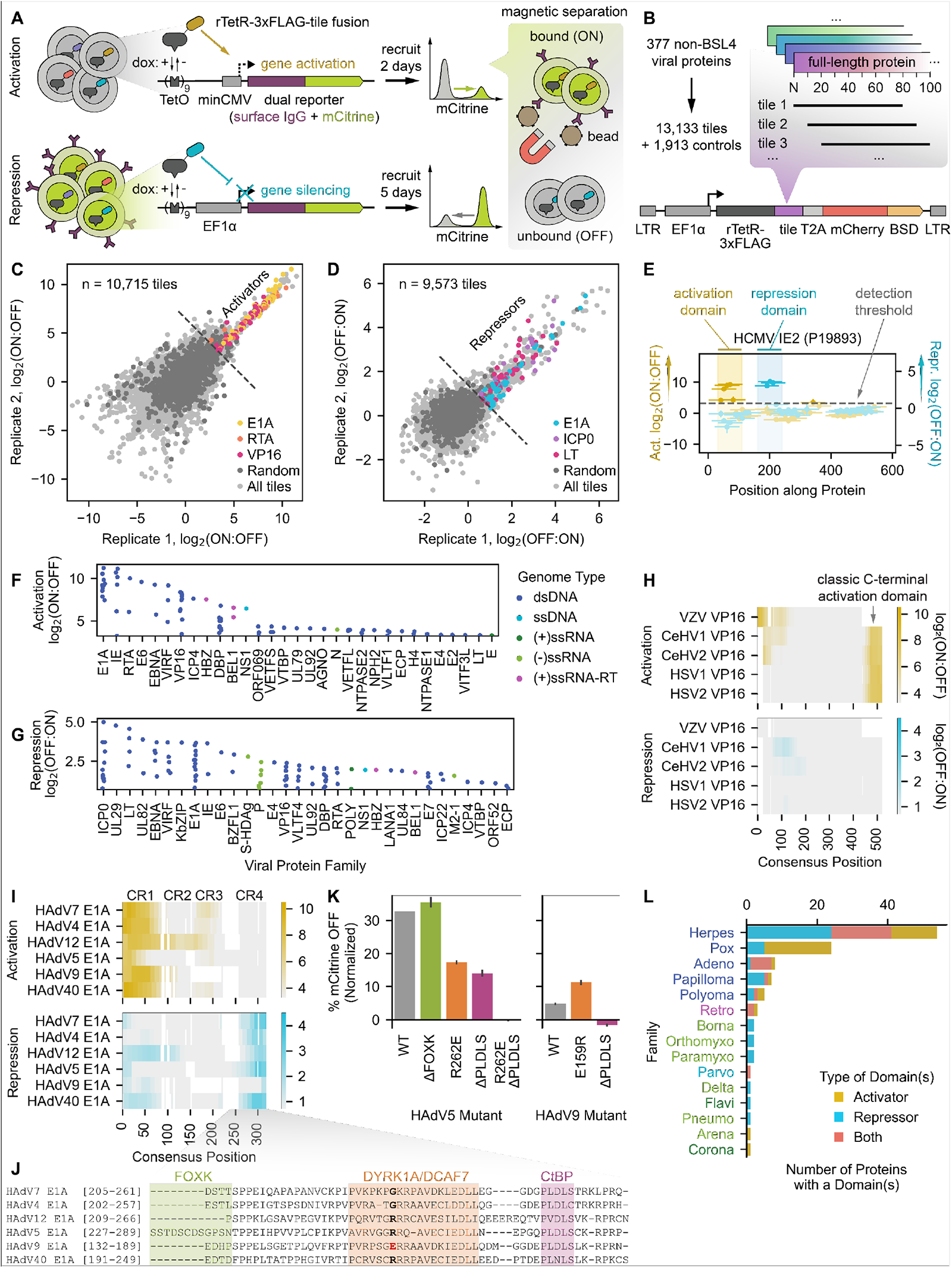
HT-recruit recovers hundreds of protein domains with transcriptional effector activity among a set of known or putative viral transcriptional regulators. **(A)** Schematic of the high-throughput recruitment (HT-recruit) approach. Library members are synthesized, cloned as fusions to the doxycycline (dox)-inducible rTetR DNA-binding domain, and delivered to cells harboring a dual reporter gene encoding both mCitrine and a surface marker that enables magnetic sorting of cells by reporter transcriptional state. Library member frequencies in the bead-bound (ON) and unbound (OFF) populations are determined by next generation sequencing to compute enrichment scores. Pooled screens are performed in cells whose reporter is under the control of a weak minimal cytomegalovirus (minCMV) promoter to measure transcriptional activation (top) or a strong EF1a promoter to measure repression (bottom). **(B)** Composition of the viral transcriptional regulator (vTR) library, which includes 80aa-long tiles sampled every 10aa for 377 proteins with known or putative transcriptional regulatory potential. BSD = blasticidin resistance gene. **(C)** Reproducibility of activation enrichment scores, log2(ON:OFF), across two replicates, with hit tiles from the well-described activators E1A, RTA, and VP16 indicated. For all analyses, the detection thresholds (dashed lines in C-G) are set as two standard deviations above the mean of the random negative controls. **(D)** Reproducibility of repression enrichment scores, log2(OFF:ON), across two replicates, with hit tiles from the repressors E1A, ICP0, and LT indicated. **(E)** Calling activation and repression domains from tiling measurements, using the HCMV IE2 protein as an example. The dashed line represents the detection threshold, and higher scores above it correspond to stronger activation (yellow) or repression (blue). Vertical spans indicate the maximum strength tile within each domain. **(F-G)** Summary of identified activation (F) and repression (G) domains, represented by their strongest tile, stratified by viral protein family, and colored by the genome type of the virus that encodes them. ds = double-stranded, ss = single-stranded, (+) = positive-sense, (-) = negative-sense, and RT = reverse-transcribed. **(H)** Multiple sequence alignment (MSA) of five herpesvirus VP16 homologs, with activation log2(ON:OFF) and repression log2(OFF:ON) enrichment scores represented as yellow and blue color mappings, respectively. **(I)** MSA of six human adenovirus (HAdV) E1A homologs with their conserved regions (CRs) indicated. **(J)** Zoomed alignment of CR4 showing known cofactor binding regions for HAdV5 E1A. A critical residue within the DYRK1A/DCAF7-binding region is bolded, with the HAdV9 mutant highlighted in red. **(K)** Quantification of the fraction of cells OFF by flow cytometry (normalized to no dox) after 5 days of recruitment of wild-type (WT) and mutant CR4 sequences from HAdV5 and HAdV9 E1A. **(L)** Summary of vTR proteins with at least one effector domain, stratified by viral family and colored by effector type. Viral family names are colored by genome type (see legend of F&G).

In order to test this method with viral proteins, we designed a library that contains strong positive controls for both activation and repression as well as proteins that have been proposed as vTRs but lack strong experimental evidence^7^. This library consists of 80aa-long protein tiles (sampled every 10aa) across 377 putative vTRs encoded by non-BSL4 human viruses^7^ as well as 80aa-long random sequences to serve as negative controls **Fig. 1B, Table S1, Methods)**. Activation and repression measurements of this library were reproducible between biological replicates **Fig. 1C&D)**. For the activation screen **(Fig. S1A)**, we computed enrichment in the ON versus the OFF population for all library members and used the scores of negative controls to define a detection threshold (**Fig. 1C, Methods)**. We identified 586 activator tiles, including those from the well-known activators E1A (from human adenovirus), RTA (from human gammaherpesviruses), and VP16 (from alphaherpesviruses) (**Fig. 1C, Table S1)**. To assess the accuracy of our assay, we performed individual recruitment experiments for a set of hits and non-hits and found good correlation between the fraction of cells ON by flow cytometry and the HT-recruit enrichment score (Spearman r = 0.86), with 95% (20 of 21) of the individually recruited hit tiles measurably activating transcription **(Fig. S1B-D)**. Similarly, we screened the same library for tiles that could repress the constitutive reporter gene **(Fig. S1E)**, defined a detection threshold of OFF versus ON enrichment scores based on the random negative control scores, and identified 476 repressor tiles, including those from the well-known repressors E1A, ICP0 (from herpesvirus), and LT (from human polyomavirus) (**Fig. 1D, Table S1)**. Screen enrichment scores correlated well with individual recruitment experiments (Spearman r = 0.92), with all 21 of the individually recruited hit tiles measurably repressing transcription **(Fig. S1F-H),** giving us confidence that our high-throughput method can reliably measure transcriptional activity.

Proteins are typically composed of structural and functional subunits called domains that are modular and can evolve independently^9^. Identifying protein domains can provide useful annotations, structural clarity, and mechanistic insight for protein and drug design purposes. One distinct advantage of screening protein tiling libraries is the ability to pinpoint the domains that are responsible for the measured function. For our assay, we defined a transcriptional effector domain as any set of two or more consecutive hit tiles or as any single hit tile positioned at the N- or C-terminus (**Fig. 1E**), and the strongest tile from each domain was used in subsequent analyses. Applying these criteria yielded 87 activation domains (**Fig. 1F**) and 106 repression domains (**Fig. 1G**) across a total of 117 proteins **(Table S3)**.

For VP16, one of the better known proteins associated with transcriptional activation and responsible for immediate early gene activation during alpha-herpesvirus infection^10^, we recovered known activation domains and, in addition, discovered previously unannotated transcriptional effector domains in some homologs (**Fig. 1H, Fig. S1I-K)**. Specifically, we detected the well-described and highly conserved tandem C-terminal activation domains present in human herpes simplex virus (HSV) 1 and HSV2 and absent in varicella zoster virus (VZV), which instead possesses a potent N-terminal activation domain **(Fig. S1J)** that shares sequence homology with part of the HSV1 and HSV2 C-terminal activation domains^11^. We also detected C-terminal activation domains in the VP16 homologs of cercopithecine herpesvirus (CeVH) 1 and CeHV2, whose natural hosts are macaque monkeys, as well as weak N-terminal activation domains that, to our knowledge, were not previously described **(Fig. S1K)**. We did not detect any activation domains in the homolog from suid herpesvirus 1, which primarily infects pigs and other non-primate animals. Interestingly, we also identified weak repression domains – some of which overlap with activation domains – within the HSV2, CeHV1, CeHV2, and SuHV1 VP16 homologs (Fig. 1H, SuHV1 data not shown, Table S3), suggesting that they may act as transcriptional repressors in certain contexts or at least engage with co-repressors.

Some of the strongest activation and repression domains we measured originate from homologs of human adenovirus (HAdV) E1A (**Fig. 1C&D, F&G)**, a highly multifunctional protein involved in cell cycle deregulation, immune evasion, and oncogenesis and known to bind over 50 cellular factors^12^. We identified effector domains in all six E1A homologs included in the vTR census (**Fig. 1I, Fig. S1L-N, representative examples)** and found that most of these domains aligned with conserved regions (CRs) previously described as having transcriptional function (**Fig. 1I**). Specifically, we identified potent transcriptional activation domains aligning with the p300-binding CR1 and the TBP/TAF-binding CR3^13^. We also identified repression domains aligning with CR4 in all homologs except HAdV9 E1A, which had a single very weak repressive tile in that region **(Fig. S1M)**.

CR4 from HAdV5 E1A, which is the best studied homolog of those in the vTR library, has been shown to contain three regions that are important for interferon response suppression and that bind the CtBP corepressor, the adaptor protein DCAF7, and FOXK transcription factors (**Fig. 1J**)^14, 15^. However, FOXK binding appears to be specific to HAdV5 E1A (**Fig. 1J**), suggesting that it is dispensable for the repressive activity we measured across homologs. Indeed, deletion of the FOXK-binding sequence had no effect on silencing (**Fig. 1K**). In contrast, mutating the DCAF7-binding region (R262E)^15^ or deleting the CtBP-binding region (PLDLS) partially reduced silencing to similar degrees, and perturbing both regions abolished silencing altogether. Consistent with these results, deletion of the CtBP-binding sequence in the weaker repressive CR4 domain from HAdV9 E1A completely abolished silencing, while installing a E159R mutation within the DCAF7-binding region to resemble the HAdV5 E1A sequence (**Fig. 1J**) increased silencing (**Fig. 1I**). These data support the observation that the combined activities of DCAF7 and CtBP are important for transcriptional repression function across E1A homologs and that the exact E1A sequence may modulate affinity for these cofactors.

Within the vTR library, we found a significant enrichment of effector domains within proteins from DNA viruses compared to RNA viruses, especially dsDNA viruses (**Fig. 1F-G, Fig S1O&P)**. This supports the observation that there is generally concordance between viral genome type and the target of encoded viral transcriptional regulators^7^. The most striking enrichment of effector domains in the vTR library was within proteins from the dsDNA herpesvirus family (**Fig. 1L**), which account for 30% of the vTR library (114 of 377) but represent 46% of proteins containing effector domains (54 of 117) (OR: 2.86, 95% CI: 1.80-4.54, Fisher’s p < 0.0001). Overall, the correlation between HT-recruit screen scores and individual flow cytometry experiments, as well as the recovery of tiles from well-described transcriptional effectors, demonstrates that our high-throughput method can quantitatively measure transcriptional activation and repression domains within viral proteins.

### Coronavirus Spike proteins harbor a conserved repression domain

Even though very few RNA virus proteins from the vTR census affected transcription, given the focus on elucidating coronavirus biology in light of the ongoing COVID-19 pandemic, we performed HT-Recruit with an unbiased library across all 280 proteins encoded by 11 bat and human coronaviruses, including SARS-CoV-2, to identify potential transcriptional activators and repressors (**Fig. 2A, Table S1)**. Both the activation and repression screens were reproducible **(Fig. S2A&B)** and their measurements correlated well with individual flow cytometry experiments **(Fig. S1D, Fig. S1H)**. While the majority of activator hit tiles barely met the detection threshold with screen scores corresponding to less than 10-15% cells ON according to our calibration curve **(Fig. S1D)**, we did identify a multi-tile activation domain within heptad repeat (HR) 2 of the MERS-CoV Spike protein (aa1191-1270) that activated the minCMV reporter in 28% of cells after two days of individual recruitment **(Fig. S1C)**. No other coronavirus Spike protein encoded an activation domain in this region, suggesting this is not a conserved function. Repressive tiles spanned a wider range of enrichment scores and constituted 47 repression domains across 36 proteins (**Fig. 2B, Table S3)**. Surprisingly, many of these repression domains mapped to the HR1 region for all 11 coronavirus Spike homologs (**Fig. 2C**). Recruitment of the strongest repressive tile from SARS-CoV-2 Spike (hereafter Spike-095) **(Fig. S2C)** greatly reduced expression of the pEF reporter gene, with 75% of cells OFF by day 5 (**Fig. 2D**).

**Fig. 2.**
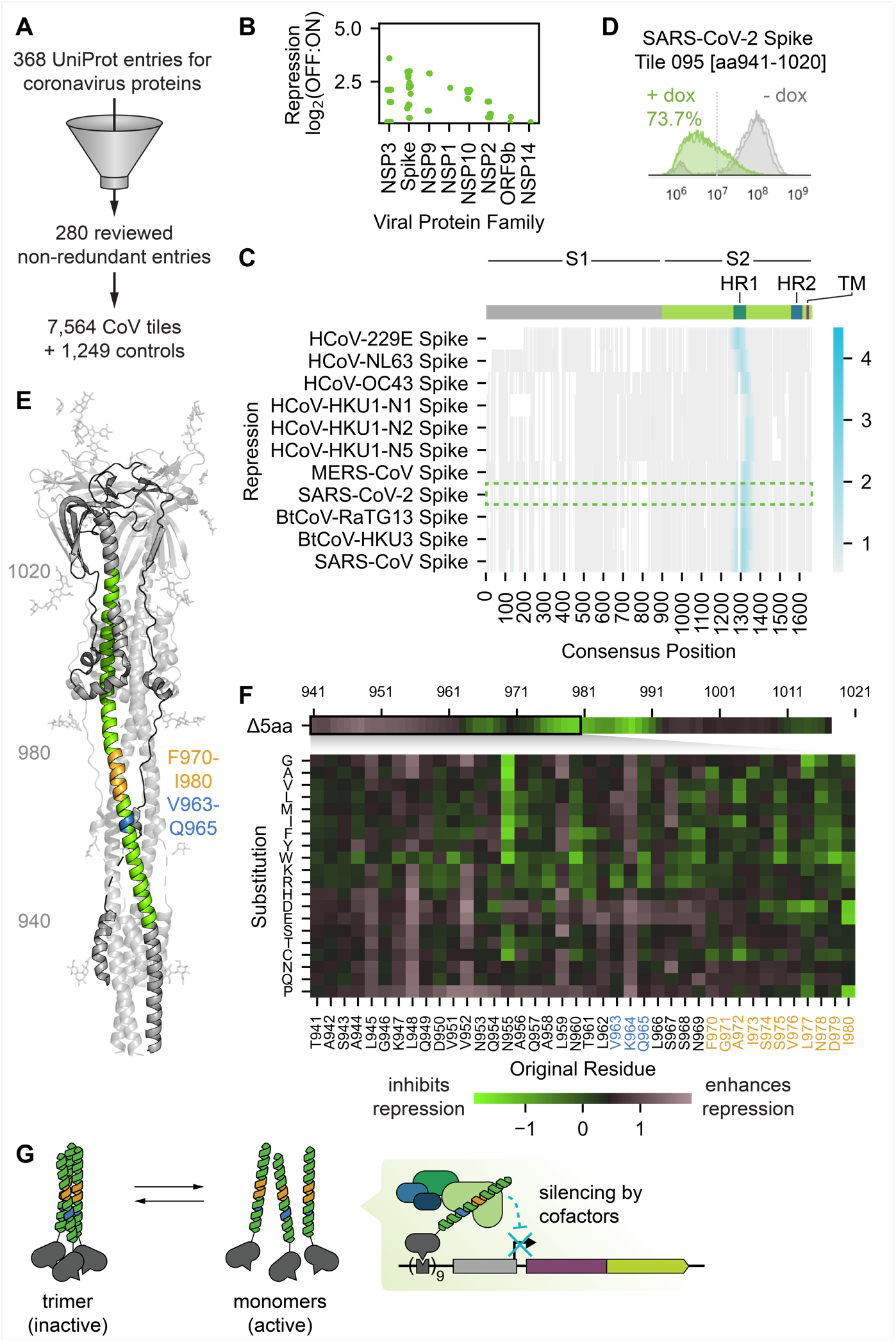
Coronavirus Spike proteins contain a functionally conserved region with transcriptional repression potential. (**A**) Coronavirus library tiling design: we tiled 280 reviewed, non-redundant entries in UniProt covering the proteomes of 11 coronaviruses to generate a library of 7,564 protein tiles supplemented with 1,249 controls. (**B**) Summary of identified repression domains, represented by their strongest tile and stratified by viral protein family. (**C**) Top: schematic of a typical coronavirus Spike protein, with the S1 and S2 fragments, heptad repeat (HR) 1 and 2 regions, and transmembrane (TM) portion indicated. Bottom: multiple sequence alignment of 11 coronavirus Spike homologs with repression domains indicated. Repression log2(OFF:ON) enrichment scores are represented as blue color mappings. The SARS-CoV-2 Spike homolog is outlined in a green box. (**D**) Flow cytometry distributions without (no dox, gray) and with (dox, green) recruitment of the SARS-CoV-2 Spike tile 095 (Spike-095), which overlaps HR1. (**E**) Structure of the SARS-CoV-2 Spike S2 fragment in the natural trimeric complex. Spike-095 is indicated in green for one monomer, with other regions of interest in the perturbation screen indicated in blue and orange. (**F**) Heatmaps of perturbation screen results. Average screen scores for overlapping deletions within the region of residues 941-1020 are shown on top. Screen scores for each substitution within residues 941-980 are shown below. (**G**) Proposed model of silencing by Spike-095 in our reporter assay.

Although we did not expect to find a transcriptional repression domain within Spike, a transmembrane protein that is best known for viral entry^16^, this domain could be liberated upon proteolysis within and escape from the endolysosome^17^ to be able to interact with chromatin regulators. To understand the physical basis for Spike-095 transcriptional repression, we designed a library comprising natural and systematically mutated Spike-095 variants and performed HT-recruit as before **(Fig. S2D&E, Table S1)**. To first assess the functional consequence of natural sequence variation within the Spike-095 region, we examined the screen scores of seven other homologs and found that the homologs from three epidemic-associated coronaviruses (SARS-CoV-2, SARS-CoV, and MERS-CoV) and a common cold coronavirus (HCoV-NL63) exhibited the strongest repression **(Fig. S2F)**. Surprisingly, these wild-type regions had comparable repression to Spike-095 while only sharing 51.3% sequence identity, akin to the amount of sequence identity that the non-repressive equivalent region from HCoV229E shares with Spike-095 (50%). This observation suggests that there is functional conservation despite sequence divergence, possibly mediated by the structure of the Spike protein.

AlphaFold predicts that Spike-095 forms a single, long alpha helix (data not shown), which is in agreement with its structure in the native Spike S2 trimer configuration that forms after entry and cleavage by host proteins (**Fig. 2E**). Thus, we hypothesized that Spike-095 could exert its transcriptional repression as either a monomer or a trimer. In order to understand which amino acids in the Spike-095 structure are important for repression, we performed the following: 1) 5aa deletion scanning with a step size of 1aa across the entire 80aa tile; 2) double and triple alanine scanning across the entire 80aa tile; and 3) a deep mutational scan of the ‘core’ region (aa 941-980) representing the intersection of all repressive tiles within the domain. Given that many mutations might only modestly affect function, which could be difficult to detect, we designed three alternatively codon-optimized sequences per mutant to improve measurement accuracy and to assess measurement precision and found that the standard deviation in screen score measurements for the three differentially coded tiles was small (mean ∼ 0.1 screen score unit) **(Fig. S2G)**. Additionally, the percent of cells OFF as measured upon individual recruitment of select library members correlated well with their screen scores **(Fig. S2H-I)**.

From our deletion scan data, we determined that the most important residues for function are in the Spike-095 core region (aa 941-980) (**Fig. 2F top, Fig. S2J)**, consistent with the intersection of the repressive tiles. Ignoring deletions that decreased repression but were non-consecutive or that sufficiently altered expression or subcellular localization of the rTetR-Spike-095 fusion **(Fig. S2J-K)**, we localized the essential region for repression to residues 977-981 (**Fig. 2E** **orange, Fig. S2K)**. Deep mutational scanning within this region revealed that biochemically dissimilar substitutions were generally detrimental to silencing, especially for normally inward-facing L977 and I980, with no substitutions enhancing activity (**Fig. 2F**). In contrast, a number of substitutions outside this region actually enhanced silencing. These included non-polar residues L945, L948, V952, or L959, which would normally contribute to stabilizing hydrophobic interactions at the Spike trimerization interface (**Fig. 2F**). Additionally, we observed enhanced silencing when almost any residue from 945-964 was substituted to proline, which is known to disrupt alpha helices like that which is predicted for the Spike-095 tile. Taken together, we propose a model in which Spike-095 may transition between its homotrimeric state to a monomeric state where residues L977 and I980 are available to interact with leucine zippers in co-repressors (**Fig. 2G**).

### Unbiased identification of activators and repressors from herpesviruses

Given their dominance in the vTR screens, we next focused on herpesviruses, which are important in human health and disease, are ubiquitous^18^, have a chromatinized dsDNA genome that persists for life^19^, and encode more proteins than most viruses^20^ (**Fig. 1L**). As such, we took a discovery-based approach to identify herpesvirus-encoded transcriptional effectors beyond those included in the vTR census, tiling nearly all known proteins (891) encoded by nine human herpesviruses and the porcine suid herpesvirus (hereafter HHV tiling library) (**Fig. 3A, Fig. S3A, Table S2)**. We found good reproducibility between replicate screens with this library **(Fig. S3B-G)**, as well as a strong correlation between individual flow cytometry experiments measuring the fraction of cells ON or OFF and the screen enrichment scores (**Fig. 3B&C)**.

**Fig. 3.**
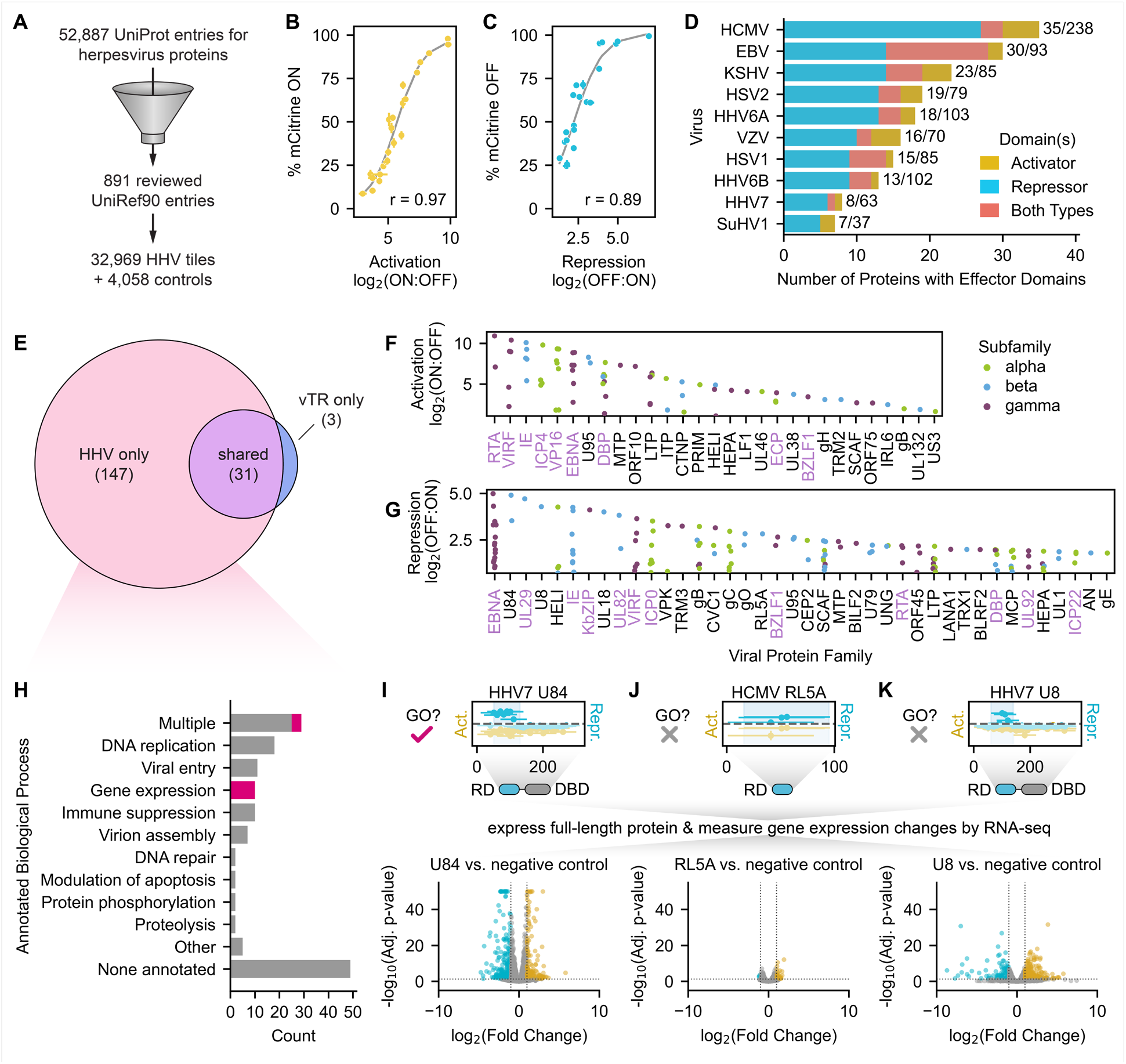
Unbiased identification of activator and repressor domains from herpesviruses. (**A**) Herpesvirus tiling library design: we compiled all 891 herpesvirus protein sequences listed in UniRef90, which collapses proteins on 90% sequence identity to limit redundancy and represent related sequences with a single high-confidence reviewed sequence, and generated a library of 32,969 tiles supplemented with 4,058 controls. (**B**) Relationship between activation log2(ON:OFF) enrichment score from the screen and the percent of mCitrine-positive (ON) cells as measured by flow cytometry after two days of recruitment for a set of individually validated tiles. Logistic fit in gray with Spearman r = 0.97. (**C**) Relationship between repression log2(OFF:ON) enrichment score from the screen and the percent of mCitrine-negative (OFF) cells as measured by flow cytometry after five days of recruitment for a set of individually validated tiles. Logistic fit in gray with Spearman r = 0.89. (**D**) Summary of herpesvirus proteins with at least one effector domain, stratified by viral species and colored by the effector domain type(s). Proteins with two or more domains of different functions or a single dual effector domain are categorized as ‘Both Types’. Bar label numerators indicate the total number of proteins with domains, and bar label denominators indicate the total number of proteins tiled for each virus (effective proteome size). **(E)** Venn diagram of all herpesvirus proteins shared between the vTR and HHV libraries in which we identified effector domains. Number of proteins per group indicated in parentheses. (**F-G**) Summary of identified activation (D) and repression (E) domains, represented by their strongest tile and stratified by viral protein family. Only the top 40 families are shown for repressors due to space constraints. Protein families labeled in lavender have at least one protein homolog for which we also measured transcriptional effector activity in the vTR screen (**H**) Summary of the biological processes associated with the 147 effector proteins uniquely identified in the HHV tiling screen. Biological process gene ontology terms associated with each protein were pulled from UniProt and assigned to high-level categories for this analysis. Proteins with gene expression-related gene ontology terms are colored in magenta. (**I-K**) Top: tiling plots from the HHV tiling screen, with a schematic showing the repression domain (RD) and the presence or absence of a predicted DNA-binding domain (DBD) for HHV7 U84 (I), HCMV RL5A (J), and HHV7 U8 (K). U84 is annotated with a gene expression-related GO term, while RL5A and U8 are not. Bottom: volcano plots from RNA-seq with expression of the full-length proteins show significantly upregulated (yellow) and downregulated (blue) genes 48 hours after induction of viral protein expression compared to a negative control expressing mCitrine instead of the viral protein.

We identified 72 activation domains and 196 repression domains across 178 proteins **(Table S3)**. Several proteins contain both types of domains (**Fig. 3D**), and sometimes activation and repression domains overlap: a subset of activator tiles spanning across all activation scores also act as weak repressors **(Fig. S3H-J, Table S2)**. Among the herpesvirus species tested, HCMV encoded the most proteins with transcriptional regulatory activity, although a higher percentage of gamma-herpesviruses EBV and KSHV proteins contain transcriptional effector domains (**Fig. 3D**).

There are 67 herpesvirus proteins that are common to the vTR and HHV tiling libraries (identical UniProt identifiers), which allows us to assess the consistency of our measurements across screens. At the tile level, we observed a strong correlation between vTR and HHV tile measurements for each of the activation **(Fig. S3K)** and repression **(Fig. S3L)** screens, with the HHV activation screen exhibiting greater sensitivity than the vTR activation screen. Additionally, 31 of the 34 (91%) herpesvirus proteins with at least one effector domain in the vTR screen also had the same effector domain in the HHV screen (**Fig. 3E**). This overlap includes well-known activators, such as VP16, RTA, and alphaherpesvirus ICP4 homologs, and repressors, such as KSHV KbZIP and alphaherpesvirus ICP0 homologs (**Fig. 3F&G)**.

Excitingly, we identified an additional 147 herpesvirus proteins unique to the HHV tiling library that possessed measurable transcriptional regulatory potential (**Fig. 3F&G),** nearly 5-fold more than the herpesvirus proteins for which we measured activity in the vTR screen. These new effectors spanned a similar range of scores **(Fig. S3M-N)** and were validated with individual flow cytometry experiments **(Fig. S3F&G)**. To better understand what was already known about these proteins and what new functional information our screen could provide, we examined the UniProt biological process (BP) GO term annotations for our hits. While two-thirds of these proteins had some annotation, only 9.5% (14 proteins) were reported to be involved in the regulation of gene expression (**Fig. 3H**) (e.g. HSV1 UL46, HHV6A IE2, HCMV UL117), with only a few of these (VP16 and ICP4 homologs) having defined effector domains in UniProt. For instance, HHV7 U84 is annotated as having a role in transcriptional regulation based on sequence homology to HCMV UL117^21^, but this activity has never been measured. Our assay identifies a strong repression domain in U84 (**Fig. 3I**), which also has a predicted DNA-binding domain^22^, suggesting a role as a viral transcription factor. Indeed, expression of full-length U84 for 48 hours produced significant changes in host gene expression profiles compared to negative control cells expressing mCitrine as measured by RNA-seq (**Fig. 3I, Fig. S3O&P, Table S5)**. Thus, for many of the proteins that do have BP GO terms related to regulating gene expression, our study provides the first experimental evidence supporting this annotation and defines the domain responsible for this activity.

The remaining effector proteins with at least one BP GO term annotation fell into several categories associated with other biological processes, including DNA replication (e.g. DNA polymerase, helicase, and DNA-binding protein homologs), viral entry (e.g. envelope glycoprotein homologs), immune suppression (e.g. HCMV UL18, EBV BLRF2, KSHV ORF52), and virion assembly (e.g. capsid assembly and tegument proteins). This finding of an additional function is consistent with the observation that viral proteins tend to be multifunctional^7, 23^.

One-third of the new transcriptional effector proteins identified in our screen (49 proteins) were not associated with any BP GO term in UniProt (**Fig. 3H**), meaning that our dataset provides the first functional annotation for these un-and under-characterized proteins. For example, the previously uncharacterized RL5A protein from HCMV harbors a moderately strong repression domain but lacks a predicted DNA-binding domain, and produces modest changes in host gene expression when expressed in its full-length form for 48 hours (**Fig. 3J, Table S5)**. Most of the differentially expressed genes are upregulated, suggesting that the repressive domain of RL5A might bind repressive cofactors and sequester them away from their target genes, leading to mild de-repression. Since it is lacking a DNA binding domain, RL5A may require additional DNA-associated factors or function in a complex with other viral proteins to exert potentially stronger transcriptional regulatory activity. In contrast, the previously uncharacterized U8 protein from HHV7 harbors a strong repression domain and a predicted DNA-binding domain, and expression of the full-length protein for 48 hours produced significant changes in host gene expression (**Fig. 3K, Table S5)**, supporting a role for this protein as a viral transcription factor. Taken together, these findings demonstrate that our high-throughput, unbiased tiling approach can discover new viral transcriptional regulators and annotate their effector domains.

### Sequence and functional comparison of EBNA family effector domains

The HHV tiling screen identified transcriptional effector domains of varying strengths (weak to very strong) within natural variants of the EBNA family proteins from EBV, enabling us to examine the functional consequences of natural sequence variation for this family. EBV is classified into two subtypes, where EBV type 1 is associated with greater prevalence and malignancy than EBV type 2 (**Fig. 4A**)^24, 25^. This typing classification is primarily driven by sequence differences between type 1 and type 2 homologs of the EBNA family proteins (**Fig. 4B**)^26^. For homologs of both types, we identified activation and repression domains in EBNA2 and EBNA3 as well as repression domains in EBNA1, EBNA4, and EBNA6 (**Fig. 4C&D**, **Fig. S4)**. These findings are consistent with previous studies that identified transcriptional corepressors and coactivators as interaction partners of the EBNA proteins^27^. For any given type 1 EBNA protein effector domain, its activation or repression strength was greater than or equal to that of its type 2 counterpart in our assay, with EBNA2 type 1 and type 2 homologs exhibiting the largest differences in activation and repression domain strengths (**Fig. 4D**).

**Fig. 4.**
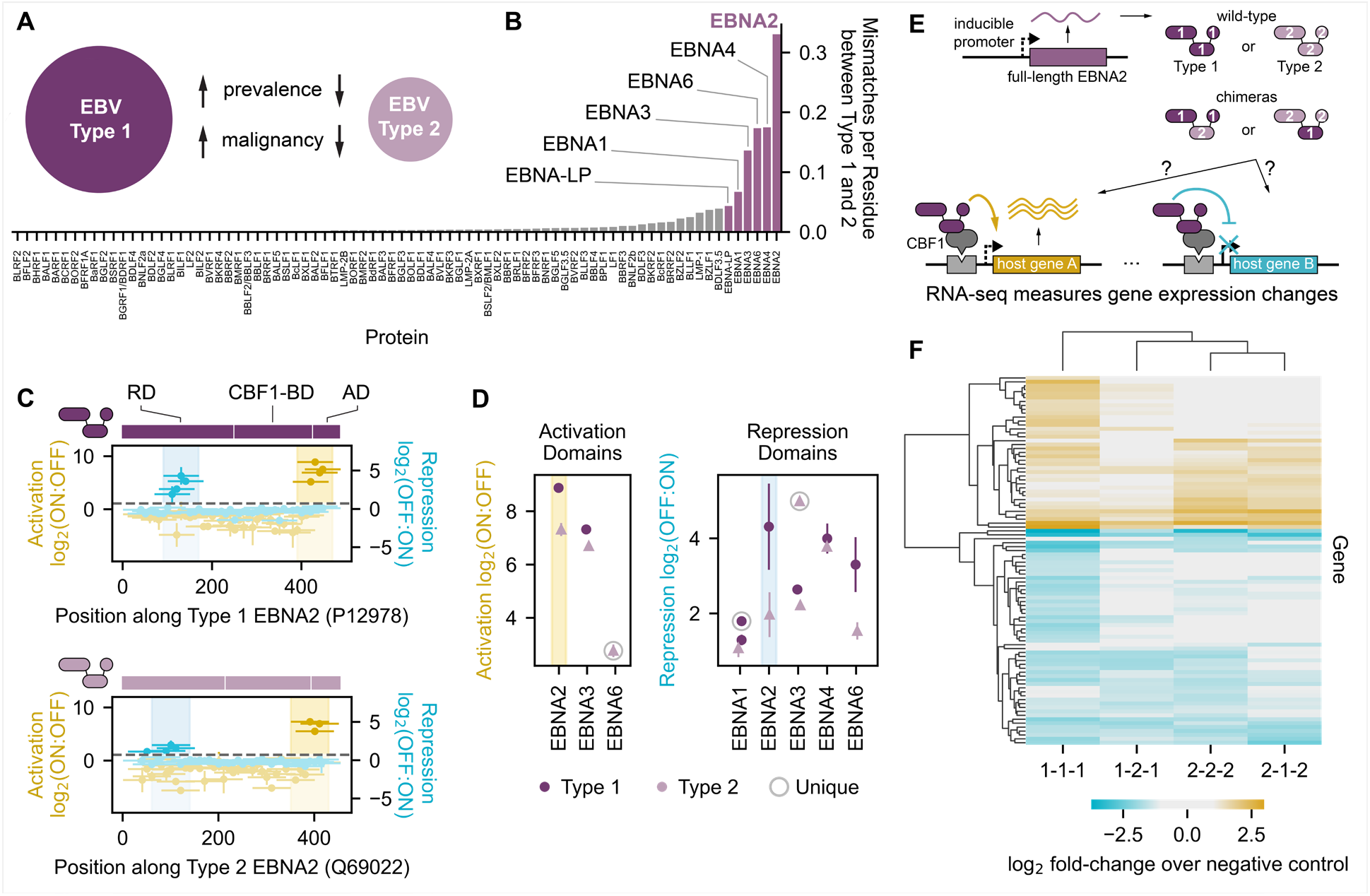
Sequence and functional comparison of EBNA family effector domains. (**A**) EBV type 1 and type 2 are associated with different clinicopathological features. (**B**) Sequence dissimilarity between type 1 and type 2 homologs of each EBV protein, quantified as the number of mismatches per residue. EBNA family proteins (highlighted) exhibit the greatest sequence dissimilarity between EBV type 1 and type 2. (**C**) Tiling plots of type 1 (top) and type 2 (bottom) EBNA2, with schematics showing domain boundaries. RD = repression domain, CBF1-BD = CBF1-binding domain (for indirect association with DNA), and AD = activation domain. (**D**) Summary of activation and repression domain strengths across type 1 (dark circles) and type 2 (light triangles) homologs of EBNA proteins. The activation and repression domains of EBNA2 are highlighted with yellow and blue spans, respectively. ‘Unique’ refers to effector domains detected in only the type 1 or type 2 homolog. (**E**) Full-length wild-type type 1 or type 2 EBNA2 (1-1-1 and 2-2-2) or chimeras with swapped effector domains (1-2-1 and 2-1-2) are expressed in K562 cells from a dox-inducible promoter (Methods) to measure differential effects on host gene expression. (**F**) Cluster map of K562 genes that are differentially expressed in at least one EBNA2 wild-type or chimera overexpression condition, as measured by RNA-seq and presented as log-2 fold changes of significantly up- or downregulated genes relative to an mCitrine-expressing negative control. Sample abbreviations reflect those shown in the schematic in (E).

Given that EBNA2 is known to associate with chromatin via its interactions with CBF1 (ubiquitously expressed) and EBF1 (not expressed in K562 cells) to affect target genes^28^, we reasoned that differences in effector domain strengths measured at a synthetic reporter gene with our tiling screen might translate to measurable differences in host gene expression upon expression of full-length EBNA2 proteins. To test this hypothesis, we expressed full-length type 1 or type 2 EBNA2 for 48 hours and harvested cells for gene expression profiling by RNA-seq and differential expression analysis by DESeq2 (**Fig. 4E, Methods, Table S5)**. To account for potential EBNA2 homolog-specific differences in binding to CBF1, we also cloned and expressed chimeras with type 1 effector domains flanking the type 2 CBF1-binding domain, and vice versa (**Fig. 4E**). For all wild-type and chimeric EBNA2 proteins, we observed significant changes in genes enriched in GO terms related to aspects of the immune response. We measured the most up- and down-regulated genes with expression of wild-type type 1 EBNA2 compared to all other proteins; in particular, type 1 EBNA2 (1-1-1) produced greater changes in expression compared to type 2 (2-2-2) (**Fig. 4F**), consistent with our tiling screen measurements of stronger type 1 effector domains (**Fig. 4D**). Interestingly, wild-type type 2 EBNA2 (2-2-2) produced more similar gene expression changes to the chimera containing the type 2 effector domains (2-1-2) than the chimera containing the type 2 CBF1-binding domain (1-2-1) (**Fig. 4F**), suggesting that these effector domains can influence genomic targets. Taken together, these data support our assay’s ability to measure differences in transcriptional effector activity between natural sequence variants both in the context of recruiting protein fragments and expressing the full-length protein.

### Sequence analyses and systematic perturbation of herpesviruses transcriptional effectors

As evident in the E1A and EBNA examples, small differences in protein sequence can produce substantial differences in transcriptional effector activity. Understanding which amino acids within transcriptional activation and repression domains are critical to and modulate function enables us to begin to understand their mechanisms of action, predict the functional consequences of viral mutations, and identify potential drug targets (**Fig. 5A**). Many eukaryotic transcriptional activation domains consist of interspersed acidic and hydrophobic residues^29–31^, while repressors fall into more categories not defined by common sequence composition^32^. In line with this, nearly all activator tiles from the HHV tiling screen have a net negative charge, with stronger activator tiles typically having greater negative charge (**Fig. 5B**). In contrast, herpesvirus repressor tiles appear to be equally likely to have net positive or negative charge (**Fig. 5B**). Both activators and repressors have an intermediate non-polar content (30-60%), and tiles with extremely low or high net charge or non-polar content generally do not exhibit effector activity (**Fig. 5B**).

**Fig. 5.**
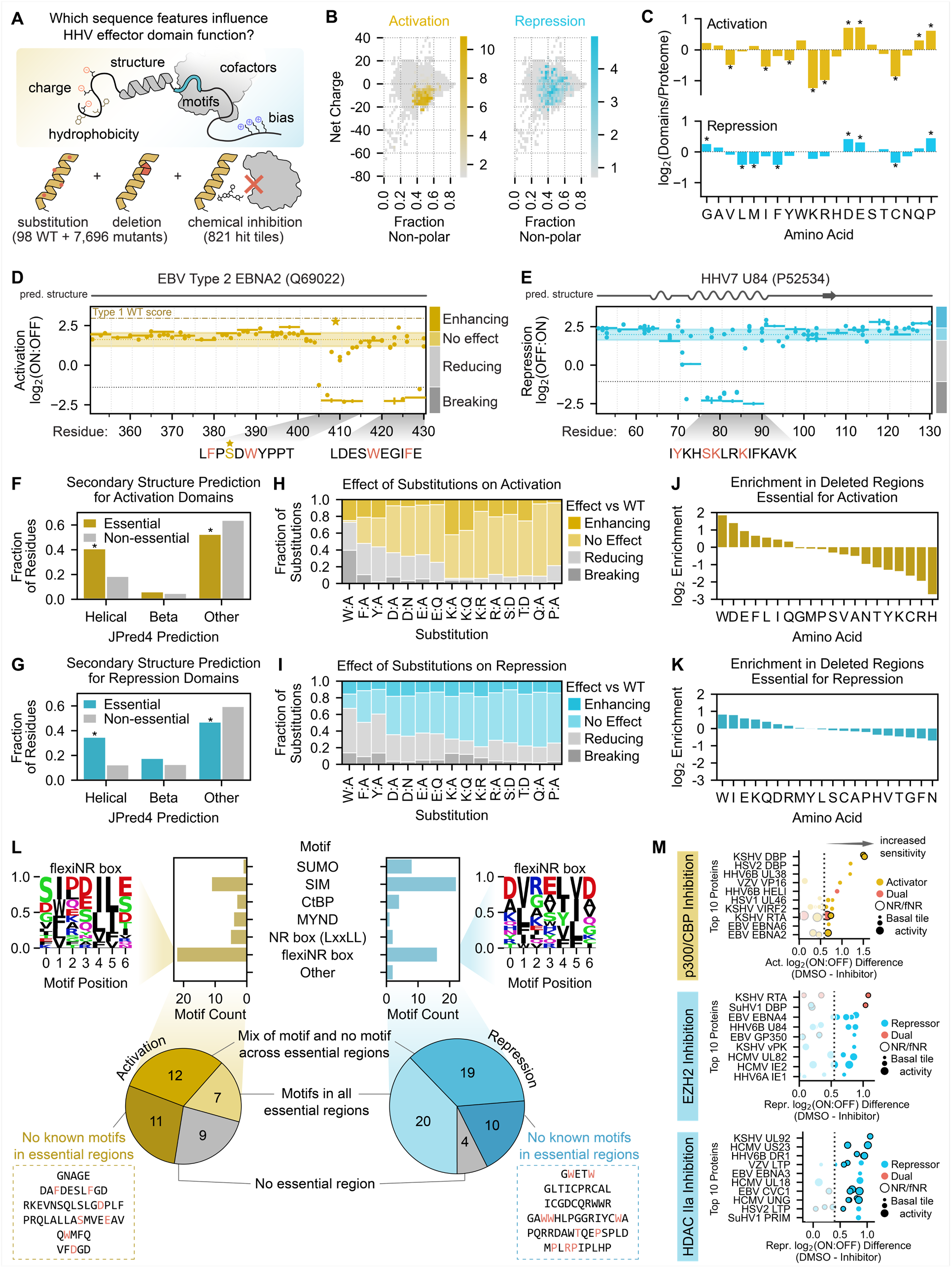
Sequence analysis and systematic perturbation of herpesvirus transcriptional effector sequences. (**A**) Overview of the sequence features examined and the perturbations performed to connect herpesvirus (HHV) effector domain sequences to their functions. (**B**) Two-dimensional histograms of net charge versus the fraction of non-polar residues for all tiles in the HHV tiling screen. Bins are colored with their maximum activation (yellow, left) or repression (blue, right) screen score. (**C**) Barplots of the log2-transformed ratios of amino acid frequencies in activation (top) or repression (domains) relative to their proteome frequencies. Positive values represent an enrichment in effector domains while negative values represent a depletion. Significant differences in amino acid frequencies were determined by the Welch’s T test and are indicated as stars. (**D, E**) Perturbation tiling plots mapping the effects of single-residue substitutions (dots) and 5aa deletions (horizontal spans) on the maximum-strength tiles from the activation domain of EBV type 2 EBNA2 (D) and repression domain of HHV7 U84 (E). JPred4-predicted secondary structures are shown above the plots, with alpha helices as squiggles, beta-sheets as arrows, and other (including unstructured) as a straight line. The shaded horizontal span represents the wild-type screen score mean plus/minus two times the estimated error (mean of all wild-type tiles shown as the yellow horizontal dotted line within) for type 2 EBNA2 (D) and U84 (E). Perturbations with scores within these regions are considered to have ‘no effect’, while those above and below are considered ‘enhancing’ and ‘reducing’, respectively. The gray horizontal dotted lines represent the detection thresholds, and thus perturbations whose scores are below this threshold are considered ‘breaking’. Deleted regions below the detection threshold are deemed essential, and their sequences are displayed below the plot, with red residues indicating single-residue substitutions that abolish activity. The brown dotted-dashed line in (D) represents the type 1 EBNA2 WT screen score for comparison of activation strength, with the yellow star representing the S409D substitution that restores type 2 EBNA2 activation to that of its type 1 counterpart. (**F-G**) Barplots comparing the fraction of residues within essential (colored) or non-essential regions (gray) regions in activation (F) or repression (G) domains that have a particular secondary structure as predicted by JPred4. (**H-I**) Effect of single-residue substitutions on activation (H) or repression (I) as measured in the perturbation screen. (**J-K**) Barplots of the log2-transformed ratios of amino acid frequencies in regions whose activity is essential to activation (J) or repression (K) relative to their proteome frequencies. (**L**) Top: counts of motifs that are enriched in essential regions. Logo of the newly proposed flexiNR box motif from all essential regions in activators (top left) or repressors (top right). Other motifs follow ELM definitions (Methods). Also shown are examples of essential sequences with no known overlapping motif (inside dashed boxes), with the residues most sensitive/critical to activity as determined by single-residue substitution in red. (**M**) Summary of the top 10 herpesvirus proteins with tiles sensitive to p300/CBP inhibition with SGC-CBP30 (top), EZH2 inhibition with tazemetostat (middle), and class IIa HDAC inhibition with TMP269 (bottom). Each dot is a tile from the viral protein indicated on the y-axis and is colored based on its effector activity. Dot size indicates the strength of the tile’s transcriptional effect in the DMSO control screen, and a black outline indicates the presence of at least one NR or flexiNR box motif (NR/fNR) in the tile. The x-axis shows the difference between screen scores in the DMSO control screen versus in the screen with inhibitor, with increasing positive values indicating increased sensitivity to the inhibitor (i.e. greater impairment of activation or repression with treatment). The dashed line in the p300/CBP inhibition screen represents the sensitivity threshold set at the mean plus two standard deviations of the repressor scores (expected to have no activity), and vice versa for the EZH2 and class IIa HDAC inhibition screens.

To better understand the sequence bases for the diverse range of transcriptional regulatory activities of herpesvirus proteins, we examined residue frequencies across effector domains. Since dual effector tiles share more sequence properties with pure activators than pure repressors **(Fig. S5A&B),** consistent with their behavior as stronger activators than repressors **(Fig. S3H)**, we grouped activator and dual effectors for all subsequent sequence analyses. Overall, activation domains are generally enriched in acidic residues and depleted in basic residues, consistent with their overall negative net charge (**Fig. 5B**), repression domains are enriched in acidic residues, and both domain types are depleted in certain non-polar residues (**Fig. 5C**). However, these enrichments do not necessarily mean that these amino-acids are important for transcriptional regulatory function of these domains.

To directly measure which residues and regions are important for transcriptional activation and repression, we systematically perturbed the amino acid sequences of the maximum-strength tiles within effector domains. In this set of high-throughput perturbation measurements, we focused on tiles that we estimated could activate or repress at least 40% of cells (**Fig. 3B&C)** so that we could measure appreciable differences in activity and test more perturbations for a smaller set of tiles. Specifically, we mutated the residues enriched in our effector domains, as well as others that have been implicated in transcriptional regulation in human cells^32^: acidic (D, E), basic (K, R), aromatic (W, F, Y), and others (S, T, Q, P). In addition, we performed deletion scanning with 5aa deletions every 5aa to identify critical regions and residues in a more unbiased manner.

Our activation and repression screens with this HHV perturbation library were reproducible **(Fig. S5C&D, S5G&H)**, individual validation experiments showed a strong correlation between percent ON or OFF and screen scores **(Fig. S5E&F, S5I&J)**, and these data identified functionally important sequences within each domain: essential regions whose deletion breaks function, as well as regions whose deletion reduces or enhances function (**Fig 5D&E, Table S4)**. For example, both EBNA2 type 1 and type 2 activation domains contain two neighboring essential regions with several phenylalanine and tryptophan residues important for activity (**Fig. 5D**). Interestingly, mutation of serine to aspartic acid at residue 409 in the weaker type 2 EBNA2 homolog restored activation levels to that of the stronger type 1 homolog (**Fig. 5D** **starred)** whose natural sequence includes an aspartic acid. Both essential regions of EBNA2 are predicted to lack secondary structure by JPred4^33^ (**Fig. 5D top**). In other examples, we identify essential sequences in regions of the protein that are predicted by JPred4 to have secondary structure. For example, our assay identified a lysine-rich essential region within HHV7 U84 harboring several critical residues whose individual substitutions were sufficient to abolish repression altogether (**Fig. 5E**). These residues mapped onto one face of a basic alpha helix that likely engages a corepressor complex (AlphaFold, data not shown). In general, essential regions within both activation and repression domains were more likely to overlap JPred4-predicted alpha-helices (**Fig. 5F&G)**, which could stabilize binding interfaces and particular side chain conformations required for activity.

Most strikingly, a high-level analysis of the functional consequences of single-residue substitutions and deletions revealed a critical role for tryptophan in transcriptional effector activity (**Fig. 5H-K**) that has not been described before. Substitution of tryptophan to alanine reduced or abolished activation and repression in 73% and 67% of cases, respectively (**Fig. 5H&I)**, consistent with the fact that tryptophan was enriched in the essential regions of both activators and repressors (**Fig. 5J&K)**. Substitution of the other aromatic residues also broke or reduced function, though less frequently (**Fig. 5H&I)**. Substitution of acidic residues reduced or abolished activation and repression in approximately 30-40% of cases, while substitution of basic residues generally only negatively affected repression and not activation (**Fig. 5I**), consistent with our findings above (**Fig. 5C**). In general, sequence bias was stronger within the essential regions of activation domains than repression domains (**Fig. 5J&K)**, most likely reflecting the greater complexity and more diverse modes of transcriptional repression also observed in human transcriptional repressors^32^.

To connect the above sequence features with how this set of effector domains might modulate transcription through the recruitment of co-repressors (CoRs) and co-activators (CoAs), we first searched for well-defined cofactor interaction motifs compiled in the ELM database and those identified in recent publications^32, 34, 35^ **(Table S6)**, ultimately focusing on those enriched in reducing/breaking regions in an initial search **(Methods)**. In essential regions of repression domains, we found several instances of SUMOylation sites, which have been connected to transcriptional repression in human cells^36^. For both activation and repression essential regions, we identified SUMO-interaction motifs (SIMs), which may bind to SUMOylated CoRs and CoAs^37, 38^ (**Fig. 5L**).

In activation domain essential regions, we found several instances of the multifunctional nuclear receptor (NR) box motif (i.e. LxxLL), which is known to engage CoAs such as p300/CBP and TFIID^39^ (**Fig. 5L**). Previous research reported instances of modified NR motifs in human proteins that can still bind their targets despite having other non-polar residues in place of leucines in the LxxLL consensus^40–42^. We also found these types of motifs, which we termed flexiNR box motifs, in the essential regions of our effector domains **(Fig. S5K, Table S6)**. With the addition of the flexiNR box motif to our list, the majority of our effector domains contain a motif for binding to a candidate cofactor: 54% of the activators and 64% of the repressors.

For 26% of activation domains and 8% of repression domains, there was no essential region whose deletion broke function (**Fig. 5L**). For nearly all of these domains, we identified two or more of the motifs listed above **(Table S6)**; thus, it is possible that upon deletion of a single motif, the other motifs may compensate to avoid total loss of activity. Conversely, for 15% of activation domains and 20% of repression domains, we identified at least one essential region but could not identify a known motif (**Fig. 5L**). While there were too few of these sequences for de novo motif finding, we found several critical acidic and aromatic residues within the activation essential regions and critical tryptophan residues within the repression essential regions, consistent with our above analysis (**Fig. 5J&K)**.

As an orthogonal approach to the identification of potential cofactors, we performed screens in the presence of chemical inhibitors of three chromatin-modifying enzymes classically associated with gene activation and silencing: SGC-CBP30, an inhibitor of the bromodomains of the CoA p300/CBP^43^ (known to bind the NR box motif); tazemetostat, an inhibitor of the histone methylation activity of the polycomb repressive complex 2 (PRC2)-associated enzyme EZH2^44^ (no known motif); and TMP269, an inhibitor of class IIa histone deacetylases (HDACs)^45^ that generally act as CoRs (no known motif). All chemical inhibition screens were reproducible, and when we compared the results of each to a DMSO-control screen, we uncovered a small set of tiles exhibiting differential activation or repression with treatment (**Fig. 5M, Table S4)**. In particular, tiles from 14 herpesvirus proteins exhibited reduced activation upon p300/CBP inhibition **(top 10 in** **Fig. 5M**). These hits include VP16, which has been shown to associate with p300/CBP^10^, as well as tiles from DBP, a family which we examine in more detail in a later section. However, sequences with NR box or flexiNR box motifs are not enriched in the set of tiles sensitive to the p300/CBP inhibition, consistent with both the ability of these motifs to recruit other CoAs and the idea that they are not the only motifs able to recruit p300/CBP. For the EZH2 and HDAC IIa inhibition screens, we identified tiles from 20 and 83 proteins, respectively, that exhibited reduced repression upon inhibitor treatment, although these changes were modest **(top 10 in** **Fig. 5M, Table S4)**. Among the effectors sensitive to EZH2 inhibition, we find sequences from the newly discovered repressor U84 (**Fig. 3I**), as well as the better studied proteins EBNA3, IE1, and IE2 (**Fig. 5M**). Surprisingly, among the tiles sensitive to HDAC IIa inhibition, we find many sequences (63%) that contain the NR or flexiNR motifs (**Fig. 5M**), suggesting these motifs recruit CoRs associated with the deacetylation pathway. These chemical screens, in conjunction with the sequence perturbations, can serve as a springboard for in-depth investigation of the molecular mechanisms associated with each effector domain.

### Investigating the importance of known and novel cofactor interaction motifs on VIRF protein functions

We identified some of the strongest herpesvirus activator, repressor, and dual effector domains in three of the KSHV viral interferon regulatory factors (VIRFs) **(Fig. S6A-C)**, which are homologous to and interact with the human IRF proteins to modulate immune signaling^46, 47^. Despite the homology between the viral and human IRF N-terminal DNA-binding domains, the effector domains of VIRF2, VIRF3, and VIRF4 differ substantially in sequence from their human counterparts (**Fig. 6A**).

**Fig. 6.**
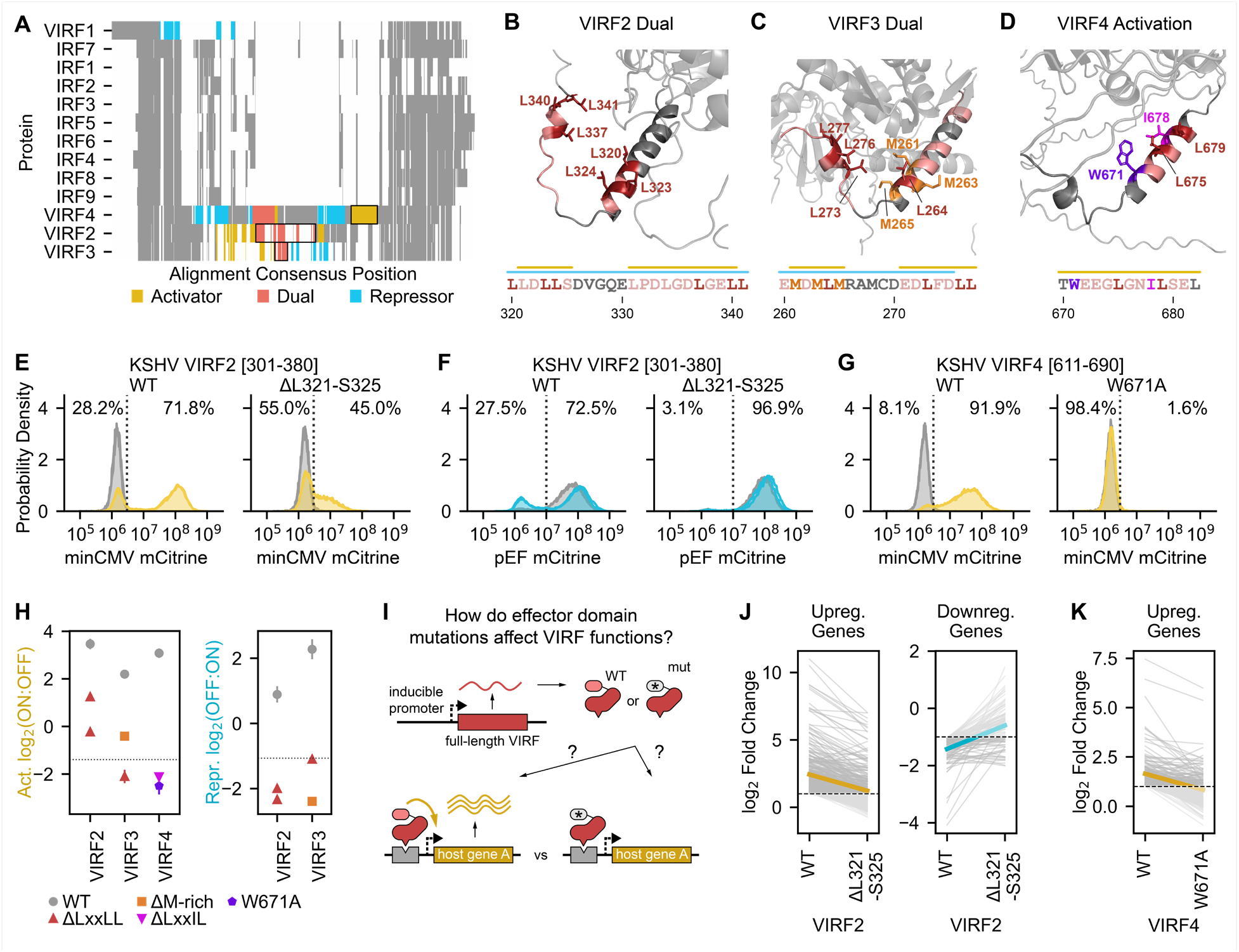
Investigating the importance of known and novel cofactor interaction motifs on VIRF protein functions. (**A**) Multiple sequence alignment of nine human interferon-regulatory factors (IRFs) and their viral homologs (VIRFs) from KSHV, showing homology across all homologs in the N-terminal DNA-binding domain and in the C-terminal region and high sequence divergence in the middle, where the VIRF2, VIRF3, and VIRF4 effector domains are located. Effector domains highlighted in subsequent panels are outlined with black boxes. (**B-D**) AlphaFold2-predicted structures highlighting portions of the dual effector domain of VIRF2 (B), the dual effector domain of VIRF3 (C), and the strong activation domain of VIRF4 (D). Residues in essential regions are colored, with key residues in the NR box (LxxLL), flexiNR box, and methionine-rich motifs in specific colors: leucines in red, methionines in orange, and isoleucine in magenta. The critical tryptophan in VIRF4 (D) is in purple. Bars above the sequences indicate regions that reduce or break activation (yellow bars) or repression (blue bars) when deleted. (**E-F**) Flow cytometry distributions of cells after recruitment for two days to a minCMV-mCitrine reporter (E) or for five days to a pEF-mCitrine reporter (F) of a wild-type or mutant (L321-S325 deletion) tile from the dual effector domain of VIRF2. The percentages of the dox-treated cells (yellow or blue) that are mCitrine-negative (OFF) and mCitrine-positive (ON) are indicated to the left and right, respectively, of the dashed line. Distributions for untreated cells are in gray. (**G**) Flow cytometry distributions of cells after recruitment for two days to a minCMV-mCitrine reporter of a wild-type or mutant (W671A) tile from the strong activation domain of VIRF4. (**H**) Summary of effector domain activity for WT VIRFs (gray) and mutants with deletions overlapping the LxxLL motif, the M-rich region, a flexiNR motif (LxxIL), or the W671A substitution. (**I**) Full-length VIRF proteins harboring either WT or mutant effector domains are expressed in K562 cells from a dox-inducible promoter (Methods) to measure differential effects on host gene expression. (**J-K**) Comparison of K562 gene expression upon overexpression of WT or mutant VIRFs, as measured by RNA-seq and presented as log-2 fold changes of significantly up- or downregulated genes relative to an mCitrine-expressing negative control. The thick yellow and blue lines indicate the mean change of all genes.

The VIRFs also differ from each other in the type and number of domains they have (**Fig. 6A, Fig. S6)** and the sequences necessary for their activity (**Fig. 6A-D**). For instance, VIRF2-4 each have a dual effector domain that activates minCMV and represses pEF (**Fig. 6A** **red, Fig. S6A-C)**. The dual effector domains of VIRF2 and VIRF3 are structurally similar (predicted alpha helices), and each contain two regions that affect their function: two NR box motifs for VIRF2 (**Fig. 6B**) and one NR box motif and a methionine-rich sequence (MDMLM) in VIRF3 (**Fig. 6C**). Deletion of the NR box motif within the VIRF3 dual effector domain completely abolishes activation and repression, while deletion of either of these two motifs in the VIRF2 dual effector domain abolishes repression but only somewhat reduces activation (**Fig. 6E-F&H, Fig. S6D-G)**. These results suggest that this motif is either bound by a single cofactor with dual transcriptional effector activities or competitively by multiple CoAs and CoRs. One candidate cofactor for the first scenario is p300/CBP, which, in addition to its well-described CoA function, can mediate transcriptional repression when SUMOylated by recruiting HDAC6^48^. Deletion of the methionine-rich region from the dual domain of VIRF3 produces a similar effect to deletion of the NR box motif: it decreases activator potential and breaks repressive function. This VIRF3 region resembles those within the EBNA2 and KbZIP repression domains **(Table S3)**, suggesting that methionine-rich sequences can coordinate CoR interactions.

VIRF4 has four effector domains (**Fig. 6A, Fig. S6C)**, none of which have been described before nor shown to interact with specific cofactors: a weak repression domain; an unstructured dual effector domain containing four critical tryptophans and several important aromatic residues **(Fig. S6H)**; a moderate-strength repression domain containing key aspartic acid and threonine residues **(Fig. S6I)**; and a strong activation domain consisting of an alpha helix with an essential tryptophan (W671) adjacent to a flexiNR box motif (LxxIL). (**Fig. 6D&G, Fig. S6J)**.

In order to understand whether the essential regions and key residues identified in our HHV perturbation screen are functionally relevant in the context of the full-length proteins, we compared the consequences of expressing full-length wild-type or mutant VIRF2 and VIRF4 on host gene expression (**Fig. 6I, Table S5)**. Overall, of the genes that are up- or down-regulated upon WT VIRF2 expression, fewer of them change significantly upon expression of an NR box mutant (deletion of residues L321-S325), and the changes are smaller (**Fig. 6J**), consistent with this mutation decreasing activation and abolishing repression (**Fig. 6E-F**). Similarly, of the genes upregulated by WT VIRF4, fewer are upregulated and to a lower extent by the W671A mutant (**Fig. 6K**), as expected from this mutation abolishing one of the VIRF4 activation domains (**Fig. 6G&H)**. These results are not due to differences in protein levels between the WT and mutant VIRFs **(Fig. S6K)**. This finding suggests that indeed the same amino acids that we identified to be important for reporter activation are also important for controlling endogenous genes in the context of the full-length protein.

### The herpesvirus DBP C-terminus regulates late gene expression and replication

Tiling across all proteins from herpesviruses identified previously unannotated, moderate-strength C-terminal transcriptional activation domains within six of the ten homologs of the herpesvirus single-stranded DNA-binding protein (DBP) **(Table S3,** **Fig. 7A-C, Fig. S7A-C)**. The DBPs are classically associated with herpesvirus genome replication, which is required for expression of late genes that encode proteins important in virion assembly^49^. Although the vTR library contained several DBPs, their inclusion in the census was due to their ability to bind single-stranded DNA rather than direct evidence for modulation of transcription.

**Fig. 7.**
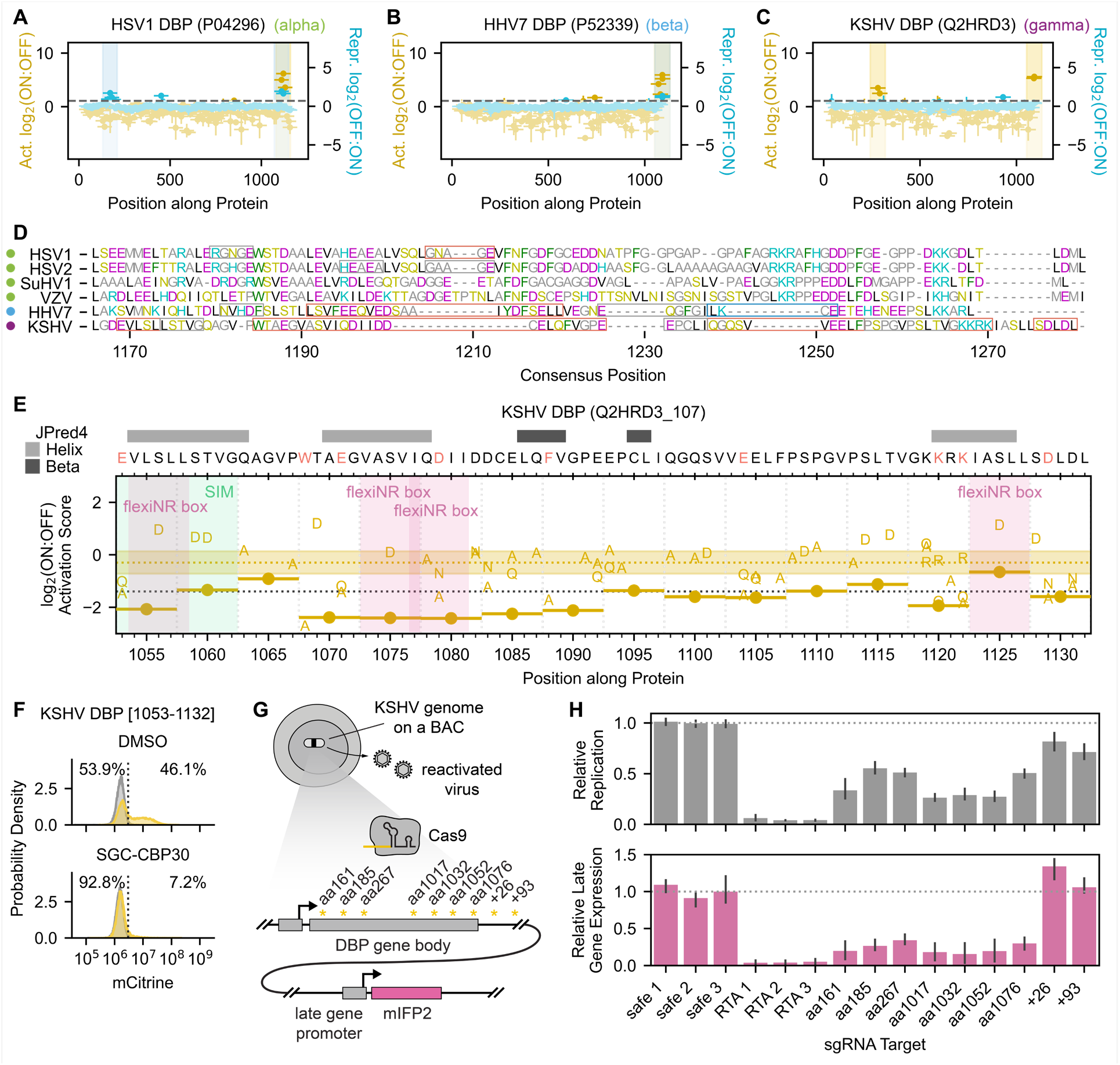
The herpesvirus DBP C-terminus regulates late gene expression and replication. **(A-C)** Tiling plots showing the C-terminal activation domain of the alpha-herpesvirus HSV1 homolog (A), beta-herpesvirus HHV7 homolog (B), and gamma KSHV homolog (C). (**D**) Multiple sequence alignment of the DBP C-terminal region for all homologs with activation potential, with amino acids colored by biochemical similarity. Colored dots to the left of the virus names indicate the herpesvirus subfamily: alpha (green), beta (blue), and gamma (purple). Red boxes indicate essential regions whose deletion breaks activity, gray boxes indicate sensitive regions whose deletion reduces activity by approximately two-fold, and the right-most blue box in HHV7 DBP indicates the SUMOylation site (LKCE) whose deletion increases activation. (**E**) Tiling plot showing the effects of 5aa deletions and single-residue substitutions on KSHV DBP activation domain activity. A SUMO-interaction motif (SIM) is highlighted with a green vertical span, and four FlexiNR box motifs are highlighted with pink vertical spans. (**F**) Flow cytometry distributions of cells with the minCMV-citrine reporter after two days of recruitment of the KSHV DBP C-terminal activation domain (aa1053-1132) with DMSO or 10uM SGC-CBP30 (p300/CBP inhibitor). (**G**) Overview of assay to measure the consequence of Cas9-induced KSHV DBP truncations on late gene expression during KSHV reactivation. Latently infected iSLK cells harbor the KSHV genome on a bacterial artificial chromosome (BAC). Targeting Cas9 to various regions of the DBP gene body produces truncated gene products whose effects on late gene expression can be measured using a KSHV genome-integrated mIFP2 reporter gene under the control of a late gene promoter. sgRNA targets are indicated by asterisks, with the approximate position indicated (residue position in gene body as ‘aa’ and base pairs past the stop codon as ‘+’). (**H**) Quantification of EdU incorporation during viral genome replication (top) and mIFP2-positivity (bottom) at 48 hours after reactivation from latency. Safe sgRNAs (safe 1-3) targeting a non-functional locus of the KSHV genome serve as negative controls that have minimal effects on DBP expression.

We identified this conserved activation domain in all four alpha-herpesvirus homologs (HSV1, HSV2, VZV, and SuHV1) (**Fig. 7A, S7A-C)**, one beta-herpesvirus homolog (HHV7) (**Fig. 7B**), and one gamma-herpesvirus homolog (KSHV) (**Fig. 7C**). We also detected mild repression potential in the same domains from HSV1, HSV2, and HHV7 (**Fig. 7A&B, Fig. S7A)**. While these domains resemble typical activation domains in that they consist of hydrophobic residues interspersed with acidic ones, the six homologs with activity do differ in sequence across alpha-, beta-, and gamma-herpesvirus subfamilies and in their essential regions (**Fig. 7D, Fig. S7D-G)**. For example, HHV7 DBP residues 1112-1116 overlap a SUMOylation site (LKCE) that we generally find in repressive domains, and its deletion strongly increases activation **(Fig. S7F-G)**. In contrast, KSHV DBP contains four flexiNR box motifs, three of which overlap essential regions (**Fig. 7E**). Moreover, tiles from the different DBP homologs show different sensitivity to the p300/CBP inhibitor, with the KSHV DBP C-terminal activation domain being the most sensitive activation domain in that screen (**Fig. 5M**, **Fig. 7F**). Taken together, we hypothesize that these C-terminal activation domains are biologically relevant and functionally conserved despite sequence and even mechanistic divergence.

Notably, deletion within the C-terminal region of the HSV1 DBP homolog has been shown to inhibit both replication and late gene expression^50^, but this has not been shown for beta- or gamma-herpesvirus DBP homologs. In order to test our hypothesis that the C-terminal activation domain of the gamma-herpesvirus KSHV DBP (residues 1053-1132) is important for late gene transcription, we used CRISPR/Cas9 to perturb this region of the KSHV genome on a bacterial artificial chromosome (BAC) in iSLK cells^51^. In this system, latently infected cells harbor an mIFP2 reporter under the control of a late gene promoter, and the expression of a dox-inducible RTA protein can reactivate these cells from latent to lytic infection (active production of KSHV virions) (**Fig. 7G**). Complete knockout of DBP in this cell culture model has been shown to prevent late gene expression as measured by a lack of mIFP2 48-hr after reactivation^51^. In this study, sgRNAs targeting positions corresponding to residues 1017, 1032, 1052, and 1076 (within one of the critical regions determined in our perturbation screen (**Fig. 7E**)) each reduced mIFP2 levels to the same degree as sgRNAs targeting the beginning of the DBP gene, indicating that deletion of this C-terminal region is functionally equivalent to complete knockout of DBP (**Fig. 7H**). EdU staining showed that viral replication was also impaired (**Fig. 7H**). Taken together, these and prior data suggest a critical, multifunctional role for the KSHV DBP C-terminus in viral genome replication and late gene expression. One model that unifies these functions would involve replication-coupled transcription, whereby DBP filament formation^52^ along the non-template strand during viral genome replication would bring p300/CBP in proximity to the newly synthesized dsDNA.

## Discussion

Viral proteins can control transcription of viral genomes and reprogram it in host cells. However, outside of a small set of viral transcriptional effector proteins that has been deeply characterized over the past several decades, most viral proteins lack functional and domain annotations supported by experimental evidence. While experimental throughput has historically been a limiting factor in gleaning this knowledge, advancements in DNA synthesis and sequencing have enabled quantitative measurements of protein functions at scale in human cells. Here, we employ high-throughput quantitative approaches to investigate transcriptional regulation across over 60,000 protein fragments across more than 1500 proteins that span the entire proteomes of 11 coronaviruses and nine human herpesviruses. Specifically, we identify the proteins that harbor activating or repressive transcriptional domains, determine where in the proteins these domains are, and the sequence features responsible for these functions. Moreover, for a subset of these proteins, we investigated the mechanistic details and consequences of these activities on host cell mRNA expression and the viral life cycle.

We first investigated a set of putative and known viral transcriptional regulators, or vTRs, to assess the efficacy of HT-recruit in recovering transcriptional effector domains. For example, we were able to identify the well-described activation domains within the HSV1 VP16 C-terminus and VZV VP16 N-terminus, as well as several effector domains that had been described for HAdV5 E1A proteins. In addition, we localized transcriptional repression activity to the N-terminus of four VP16 homologs. To our knowledge, this study is the first to directly compare the strengths of multiple VP16 and E1A effector domains across homologs. Overall, our assay identified transcriptional regulatory domains in over one hundred proteins included in the vTR census (117/377). While all vTR members have some evidence supporting their inclusion in the census (such as DNA or RNA binding), it is possible that members of the vTR census in which we do not identify any effector domain with our method either: 1) have effector domains where the necessary sequence is larger than 80aa, 2) require other viral or human cofactors that are not present in our cells, or 3) bind DNA but do not contain transcriptional effector domains, and enact their function in cells by competing with human transcription factors for binding across the genome. The approaches developed here can be further extended to address these questions. In addition, the vTR library contains a mixture of proteins that may act on DNA and/or RNA substrates, yet we only measured the ability to affect transcription from a dsDNA template. Indeed, the authors of the vTR census show that the viral genome type that encodes a protein is generally concordant with the protein’s particular substrate^7^. This agrees with the enrichment of transcriptional effector domains identified in dsDNA virus proteins and the relative dearth identified in RNA virus proteins in our reporter assays. A similar high-throughput approach using RNA fluorescent reporters could be used to measure the activities of RNA viral regulators.

In our unbiased screens for coronavirus-encoded transcriptional regulators, we identified relatively few hits, which, as discussed above, is unsurprising for a family of RNA viruses. However, we found that all 11 Spike homologs tiled in this library harbored a repression domain mapping to HR1. Deletion and deep mutation scanning of Spike-095 localized repression activity to the hydrophobic residues within a small region (aa977-981) that resembles a leucine zipper. Leucine zippers are common as protein dimerization domains and would allow Spike-095 to interact with leucine zipper-containing host factors involved in transcriptional repression, although such cofactors remain to be elucidated. Interestingly, Spike-095 has moderate homology to other proteins that contain leucine zipper domains, including MAX, which is known to interact with a number of transcriptional regulatory proteins and complexes, suggesting possible mimicry^53^. In the native context, the Spike-095 region appears to stabilize trimers of Spike S2 fragments through hydrophobic interactions at the trimerization interface, which includes the key hydrophobic residues within aa977-981. However, nearly all mutations upstream of this critical region that would impair trimerization and destabilize the helical structure actually enhanced repression, and vice versa. So, while the rTetR-Spike-095 fusions likely trimerize, interactions with host factors may only be possible in the monomeric form when aa977-981 are exposed. Such a model where competing homo- and hetero-typic interactions modulate function may be useful to synthetic biologists developing sensors and other tools.

These findings of a transcriptional repression domain in Spike homologs are unexpected given the classical role of viral transmembrane glycoproteins as critical factors in tropism, fusion, and entry^54^. In SARS-CoV-2, HR1 is normally sheathed by the Spike S1 portion, which is shed after a series of interactions and cleavage events by host proteases on the cell surface or in the endolysosomal pathway. Analysis of potential Spike cleavage sites by human proteases has identified several possible cathepsin-sensitive sites flanking the regions explored in this study^55^, with cathepsin L shown to cleave Spike^56^. This type of cleavage could create Spike fragments more similar to the one that we detect as a repressor. Moreover, recently, the Spike protein has been detected in the nucleus, albeit at low frequency^57^. However, there has not been prior evidence for Spike being involved in transcriptional regulation. Therefore, it remains to be determined whether our findings about small Spike protein fragments are physiologically relevant upon coronavirus infection.

Our unbiased screens for herpesvirus-encoded transcriptional effectors identified activation or repression domains in 178 of the 891 proteins (20%) included in the tiling library, a surprisingly high hit rate. At the most basic level, viruses need to enter a host cell, replicate their genome, and produce structural components to package these genome copies. Only a small set of genes is required for these processes (e.g. five total for rabies virus), yet herpesviruses encode 80-200 genes each, with many of these genes harboring transcriptional effector domains as measured in our assay. This finding suggests that together these proteins may enable more complex regulation of viral and host gene expression. This may be reflected in the unique features of herpesviruses, such as their near universal prevalence, lifelong infection with repeated transitions between active and inactive states, their immune evasiveness, and their implication in chronic, autoimmune, and neurodegenerative diseases^58–60^. Some of the strongest effectors we identified were late gene proteins and latency factors from gamma-herpesviruses, which can infect many cells but establish latency in B cells. It is possible that our K562 model cells, which are also derived from bone marrow, express similar transcriptional and chromatin cofactors. Follow-up studies that screen herpesvirus proteins in different cell types will provide mechanistic insight into cell-type specific consequences of infection.

It is also important to understand how the activities of isolated effector domains connect to the function of the full-length protein, especially in models relevant to viral infection. Encouragingly, we did find a positive correlation between effector strength and the likelihood that the full-length protein contains a predicted nuclear localization signal **(Fig. S3Q&R)**, suggesting that many of these proteins are likely to act in the nucleus and affect host gene expression. Moreover, when we expressed individual full-length proteins that contain predicted or known DNA binding domains, we measured significant changes in gene host expression (e.g. HHV7 U8 and U84 in Fig. 3I, VIRFs in Fig. 6J&K), and these changes were affected by mutations that impaired effector domain function (VIRFs in Fig. 6J&K). Finally, for one member of the newly discovered family of activation domains from the herpesvirus DBPs, we tested its role within the context of the full-length protein and a KSHV viral model, and showed that deletions of this domain impair viral replication and late-gene expression. We hope this large quantitative dataset will massively expand herpesvirus protein annotations, which are largely lacking, and that similar integrative approaches with full-length viral proteins and mutants that lack transcriptional activity will help virologists interpret and build upon these findings.

Sequence analyses and perturbations revealed that herpesvirus activators and repressors share some properties with human ones, which is unsurprising given that they must work with host machinery. One key finding unique to herpesvirus effectors is the importance of tryptophan residues to both activation and repression: 73% and 67%, respectively, of all single substitutions reduced or completely abolished activity. These critical tryptophans tend to be surrounded by acidic residues and some hydrophobic residues for many effectors, suggesting that there may be a way to predict critical regions of activity and determine whether these rules extend to other dsDNA viruses. Related to this trend, we identified several variations of the multifunctional NR box motif (LxxLL) within essential regions of herpesvirus activation and repression domains, as well as essential regions with no known motif. Future studies using wild-type and mutant proteins lacking these essential regions can elucidate the interaction partners associated with these novel motifs and responsible for these effector activities. More broadly, understanding the rules that underlie activation and repression would enable protein engineers to design novel, synthetic transcriptional effectors. In the meantime, this study provides a rich repertoire of short activator, repressor, and dual effector domains spanning a range of strengths and acting through a variety of cofactors that should expand and improve the synthetic gene regulation toolkit.

The inclusion of homologs from different viral species and strains allow us to appreciate 1) how function can be conserved despite natural sequence variation (e.g. DBP activation domain) as well as 2) how homolog-specific functions can arise despite high sequence similarity (e.g. EBNA2 repression domain). The ability to design and simultaneously test the functional consequences of thousands of deletions and substitutions allows us to map essential regions of activity for hundreds of effector domains, something that has not been possible in systems with live virus. Additional screens with chemical inhibition of chromatin-modifying enzymes and investigation of host gene expression changes in the presence of individually expressed viral proteins help further elucidate the mechanisms and consequences of these transcriptional regulatory activities. Thus, these high-throughput, quantitative synthetic biology approaches provide a powerful way to understand the physical basis for viral protein function and complement traditional virological methods, with the added benefit of enabling investigation of proteins from viruses that otherwise cannot be easily grown in cell culture. This knowledge will facilitate *in silico* drug screening and the development of new antivirals and vaccines. In summary, our catalog of viral protein sequences that act as transcriptional effectors in human cells, together with their functional mutants, can serve as a resource for interpreting viral protein function and sourcing components for synthetic biology tools.

## Supplemental Figures

**Fig. S1.**
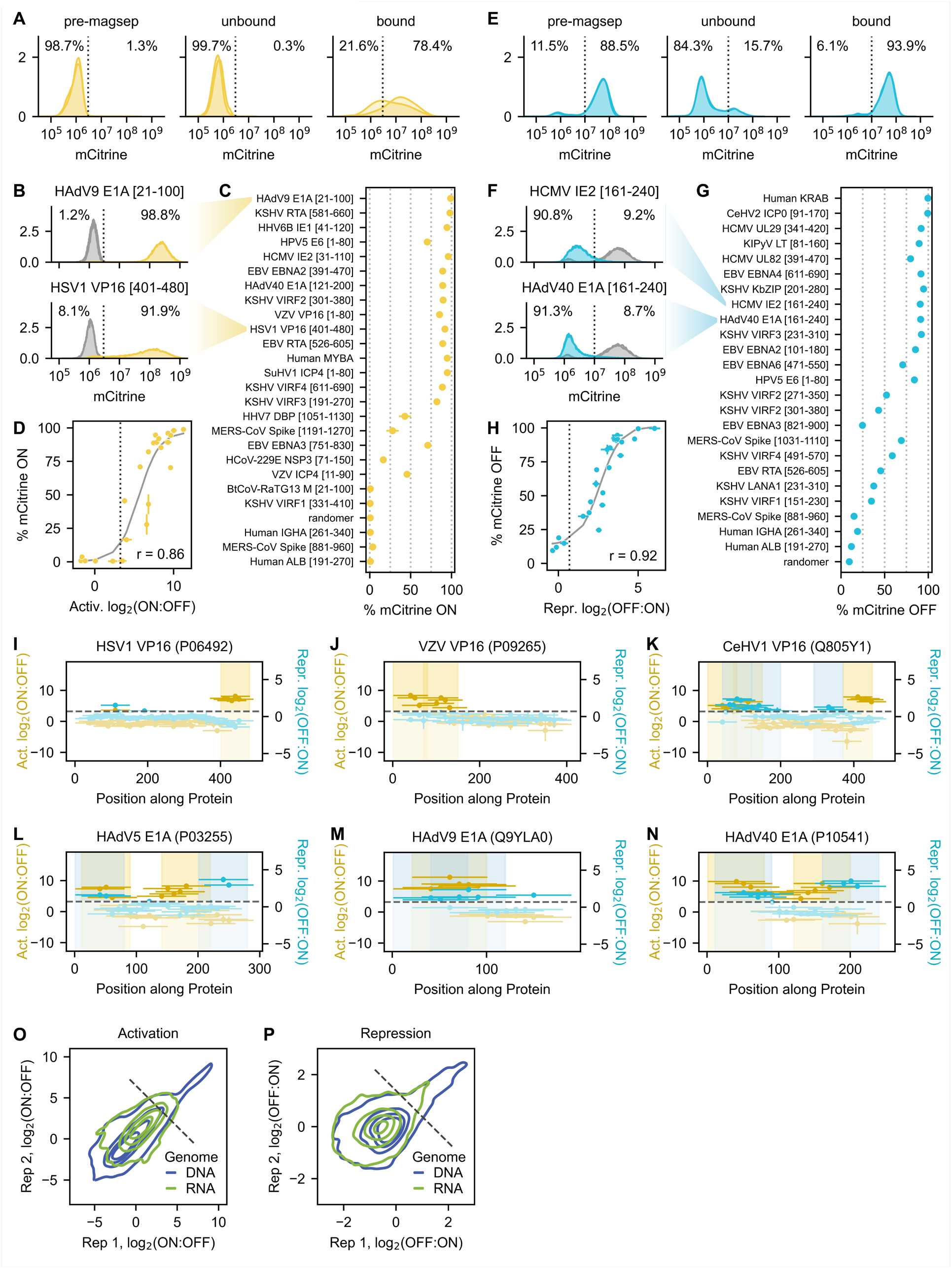
vTR and CoV activation screen details and validation. (**A**) Flow cytometry distributions on day two of recruitment for replicate screen minCMV reporter cells before and after magnetic separation. (**B**) Example flow cytometry distributions on day two of recruitment for individually validated activator tiles from human adenovirus 9 (HAdV9) E1A and herpes simplex virus 1 (HSV1) VP16. The percentages of the dox-treated cells that are mCitrine-negative (OFF) and mCitrine-positive (ON) are indicated to the left and right, respectively, of the dashed line. (**C**) Summary of activation strengths of individually validated tiles as assessed by flow cytometry. Tiles are ranked by screen score, with percent of ON cells shown on the x-axis. (**D**) Relationship between activation log2(ON:OFF) enrichment score from the screen and the percent of ON cells for the set of individually validated tiles in (C). Logistic fit in gray with Spearman r = 0.86. The gray dashed line at x = 3.25 represents the detection threshold. (**E**) Flow cytometry distributions on day five of recruitment for replicate screen pEF reporter cells before and after magnetic separation. (**F**) Example flow cytometry distributions on day five of recruitment for individually validated repressor tiles from human cytomegalovirus (HCMV) IE2 and HAdV40 E1A. (**G**) Summary of repression strengths of individually validated tiles as assessed by flow cytometry. Tiles are ranked by screen score, with percent of OFF cells shown on the x-axis. (**H**) Relationship between repression log2(OFF:ON) enrichment score from the screen and the percent of OFF cells for the set of individually validated tiles in (G). Logistic fit in gray with Spearman r = 0.92. The gray dashed line at x = 0.69 represents the detection threshold. (**I-K**) Tiling plots for three VP16 homologs included in the vTR library: herpes simplex virus 1 (HSV1) VP16 (I), HSV2 VP16 (J), and varicella zoster virus (VZV) VP16 (K). Activation and repression domains are highlighted as vertical spans. (**L-N**) Tiling plots for three E1A homologs included in the vTR library: HAdV5 E1A (L), HAdV9 E1A (M), HAdV40 (N). (**O**) Activation screen reproducibility plot as in Fig. 1C, but rendered as contours and colored by virus genome type (DNA or RNA). (**P**) Repression screen reproducibility plot as in Fig. 1D, but rendered as contours and colored by virus genome type.

**Fig. S2.**
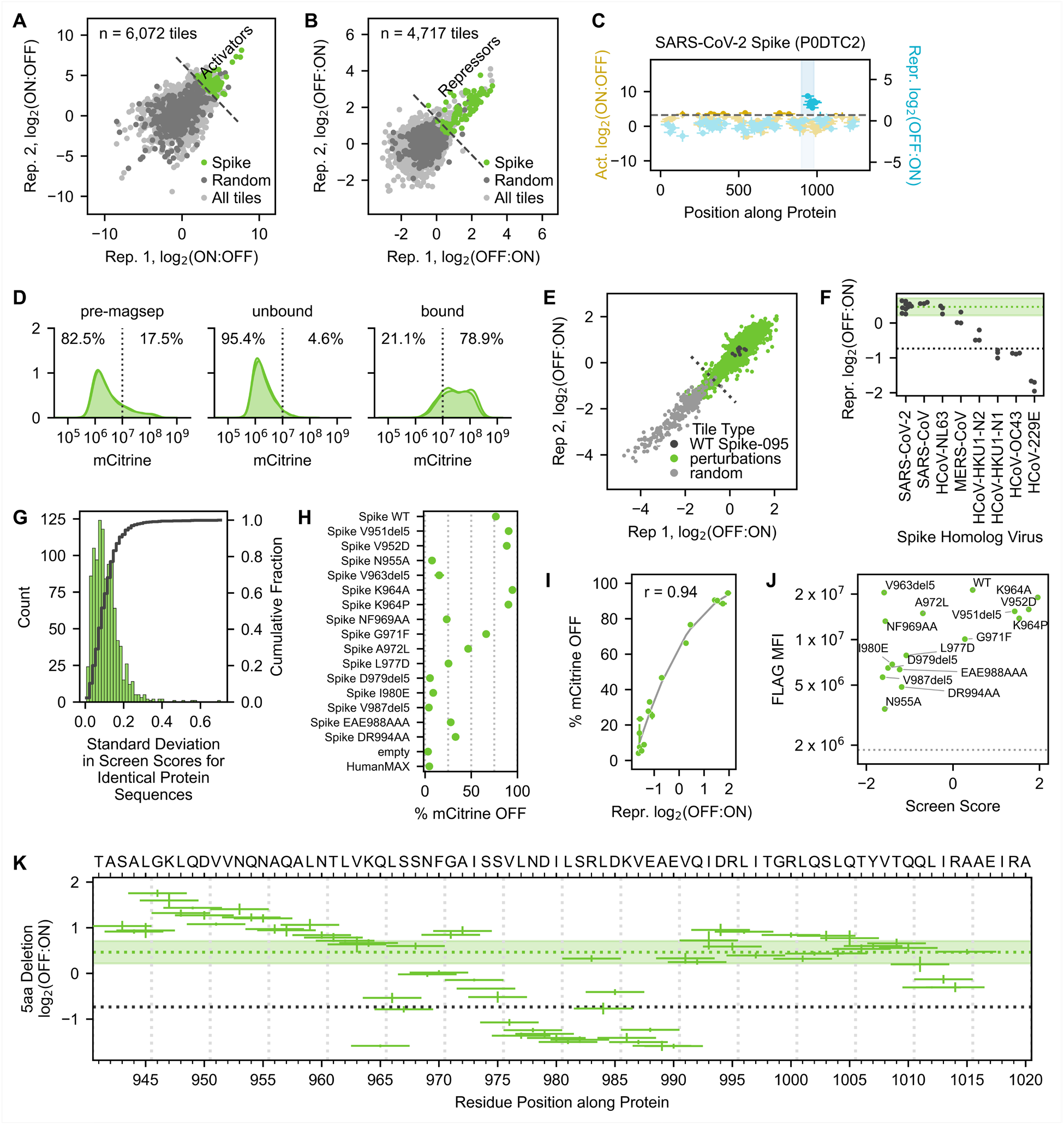
SARS-CoV-2 Spike perturbation screen details and validation. (**A-B**) Coronavirus screen reproducibility plots for activation (A) and repression (B), with tiles from Spike protein homologs indicated. (**C**) Tiling plot of SARS-CoV-2, highlighting the maximum-strength tile (Spike-095) from the repression domain overlapping the heptad repeat 2 (HR2) region. (**D**) Flow cytometry distributions on day seven of recruitment for replicate screen pEF reporter cells before and after magnetic separation. (**E**) Spike-095 perturbation screen reproducibility plot, showing repression log2(OFF:ON) enrichment scores across two replicates. The detection threshold (dashed line) is set as two standard deviations above the mean of the random negative controls. **(F)** Screen scores for wild-type sequences of the SARS-CoV-2 Spike tile 095 (aa941-1020) and the corresponding sequences from seven other coronavirus Spike homologs. (**G**) Distribution of standard deviations in screen scores between alternatively coded library tiles (i.e. identical protein sequence but different DNA sequences). (**H**) Summary of repression strengths of individually validated tiles as assessed by flow cytometry. Tiles are ranked by screen score, with percent of mCitrine-negative (OFF) cells shown on the y-axis. (**I**) Relationship between repression log2(OFF:ON) enrichment score from the screen and the percent of OFF cells for the set of individually validated tiles in (H). Logistic fit in gray with Spearman r = 0.94. (**J**) Fusion protein levels (rTetR-3xFLAG-effector) are estimated using anti-FLAG staining and flow cytometry. Scatterplot of screen score against FLAG MFI (proxy for protein levels). Dashed line represents background FLAG staining signal from WT cells (no protein expression), showing all mutants are expressed, regardless of their screen score. (**K**) Perturbation plot showing the effect of 5aa deletions on transcriptional repression by Spike-095, with the 5aa region deleted represented as a horizontal bar. The green dashed line and solid horizontal span represents the mean of the ten alternatively coded wild-type Spike-095 sequences plus and minus two standard deviations. The black dashed line at y = -0.73 represents the detection threshold.

**Fig. S3.**
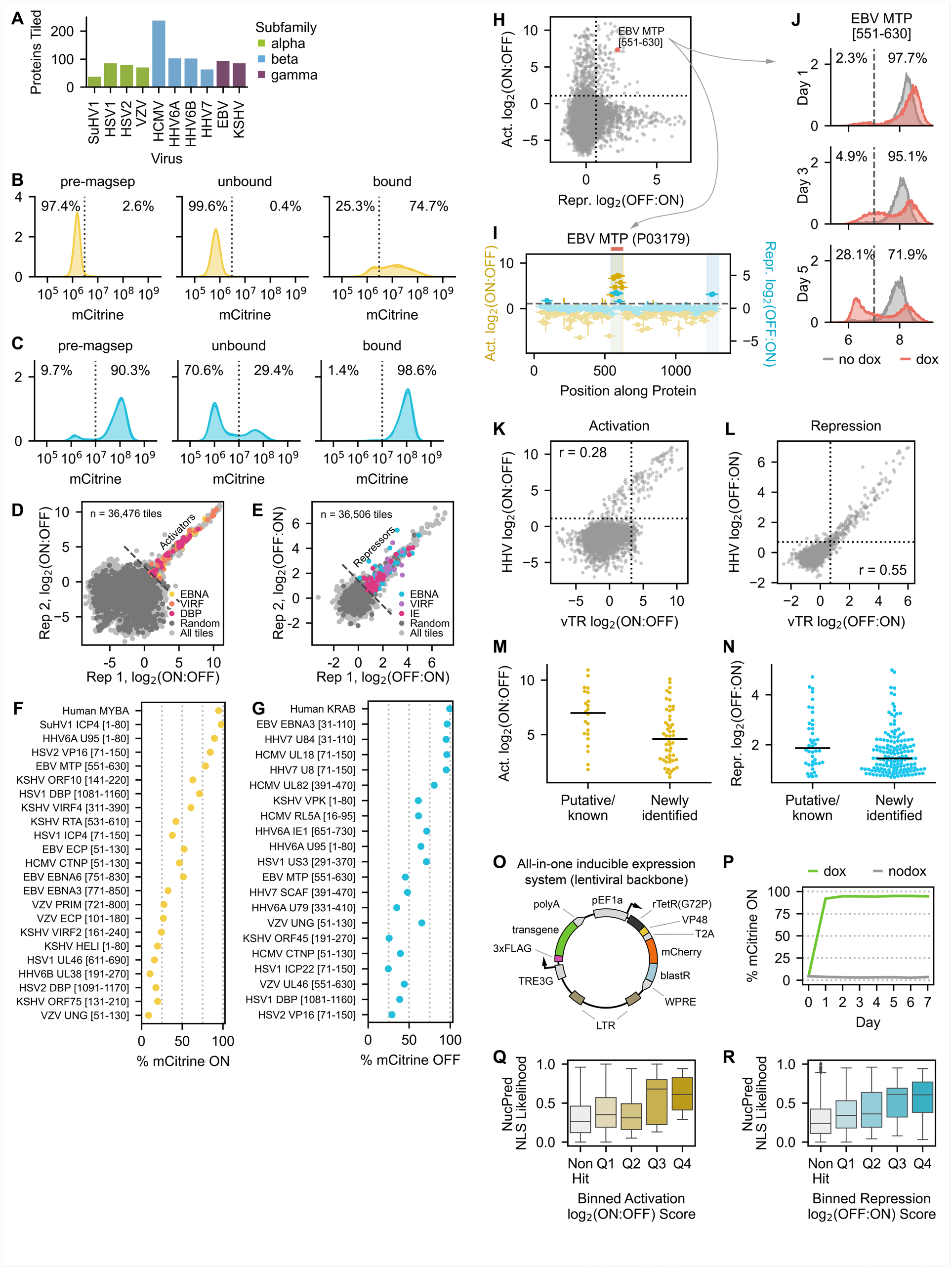
Herpesvirus tiling screen details and validations. (**A**) Summary of the number of proteins tiled for each herpesvirus species, colored by subfamily. Tiled proteins were pulled from UniRef90 and represent nearly complete proteome coverage. (**B-C**) Flow cytometry distributions on day two of recruitment for replicate screen minCMV reporter cells (B) and on day five of recruitment for replicate screen pEF reporter cells (C) before and after magnetic separation. (**D**) Reproducibility of activation log2(ON:OFF) enrichment scores across two replicates, with hit tiles from EBNA, VIRF, and DBP proteins (investigated later) indicated. The gray dashed line represents the detection threshold, which is 1.08. (**E**) Reproducibility of repression log2(OFF:ON) enrichment scores across two replicates, with hit tiles from EBNA, VIRF, and IE proteins (former two investigated later) indicated. The gray dashed line represents the detection threshold, which is 0.70. (**F**) Summary of activation strengths of individually validated tiles as assessed by flow cytometry. Tiles are ranked by screen score, with percent of mCitrine-positive (ON) cells shown on the y-axis. (**G**) Summary of repression strengths of individually validated tiles as assessed by flow cytometry. Tiles are ranked by screen score, with percent of mCitrine-negative (OFF) cells shown on the y-axis. (**H**) Activation versus repression screen scores. Tiles with dual activities are in the top right quadrant. One example from EBV MTP is highlighted in red. (**I**) Tiling plot of EBV MTP with its dual effector domain highlighted with a red bar. This domain displays strong activation and moderate repression potential. (**J**) Gene silencing dynamics of the dual effector tile from EBV MTP highlighted in (H) upon recruitment at the pEF reporter: mCitrine levels initially increase in all cells, then subsequently bifurcate, with one population maintaining elevated mCitrine levels relative to the no dox sample and the other silencing completely by day 5 of recruitment. These dynamics are observed for several validated dual effector tiles (data not shown) and are similar to those observed for a set of dual effectors tiles from human transcription factors, albeit at a different promoter^32^. (**K-L**) Scatterplot of the average activation (K) or repression (L) screen scores for tiles shared between the vTR and herpesvirus (HHV) libraries. (**M-N**) Comparison of activation (M) and repression (N) domain strengths for putative/known herpesvirus effector proteins present in both the vTR and HHV tiling screens versus those newly identified in the HHV tiling screen. (**O**) Plasmid design for all-in-one inducible expression of full-length viral proteins on a third generation lentiviral backbone. The pEF1a promoter constitutively expresses a non-leaky reverse tetracycline repressor (rTetR(G72P) mutant) fused to the strong activation VP48 domain, which, upon doxycycline (dox) addition, can activate the TRE3G promoter for expression of a transgene: either a full-length viral protein or mCitrine as a negative control. The rTetR(G72P)-VP48 is linked by a ribosome-skipping T2A sequence to a fusion protein of mCherry and an enzyme conferring resistance to blasticidin (blastR) for visualization and selection purposes. (**P**) Time course showing sustained, robust expression of an mCitrine transgene with the system described in (O) with continued doxycycline treatment. (**Q-R**) Boxplots of predicted NLS (NucPred score) across full-length viral proteins, binned by activation (Q) or repression (R) screen score of their effector domains. Q1 through Q4 represent the first through fourth quartiles of effector strength.

**Fig. S4.**
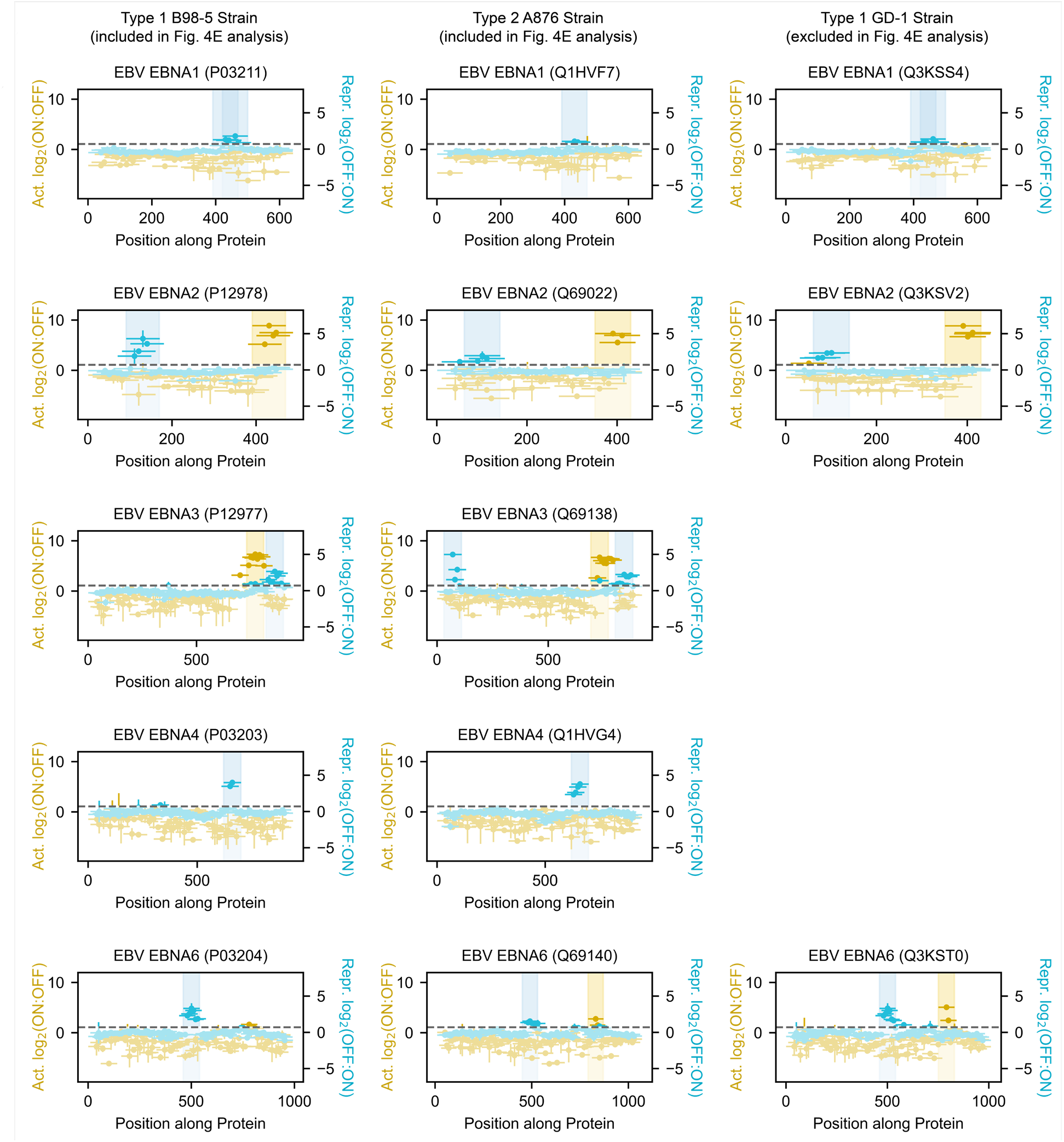
Effector domains across EBNA protein homologs. Tiling plots for all EBNA proteins with effector domains identified in the HHV tiling library. Rows delineate EBNA protein types, while columns delineate EBV strains. EBV type 1 and type 2 are represented by the classical B95-8 and A876 strains, respectively, which were used for the analysis in Fig. 4D and subsequent RNA-seq experiments.

**Fig. S5.**
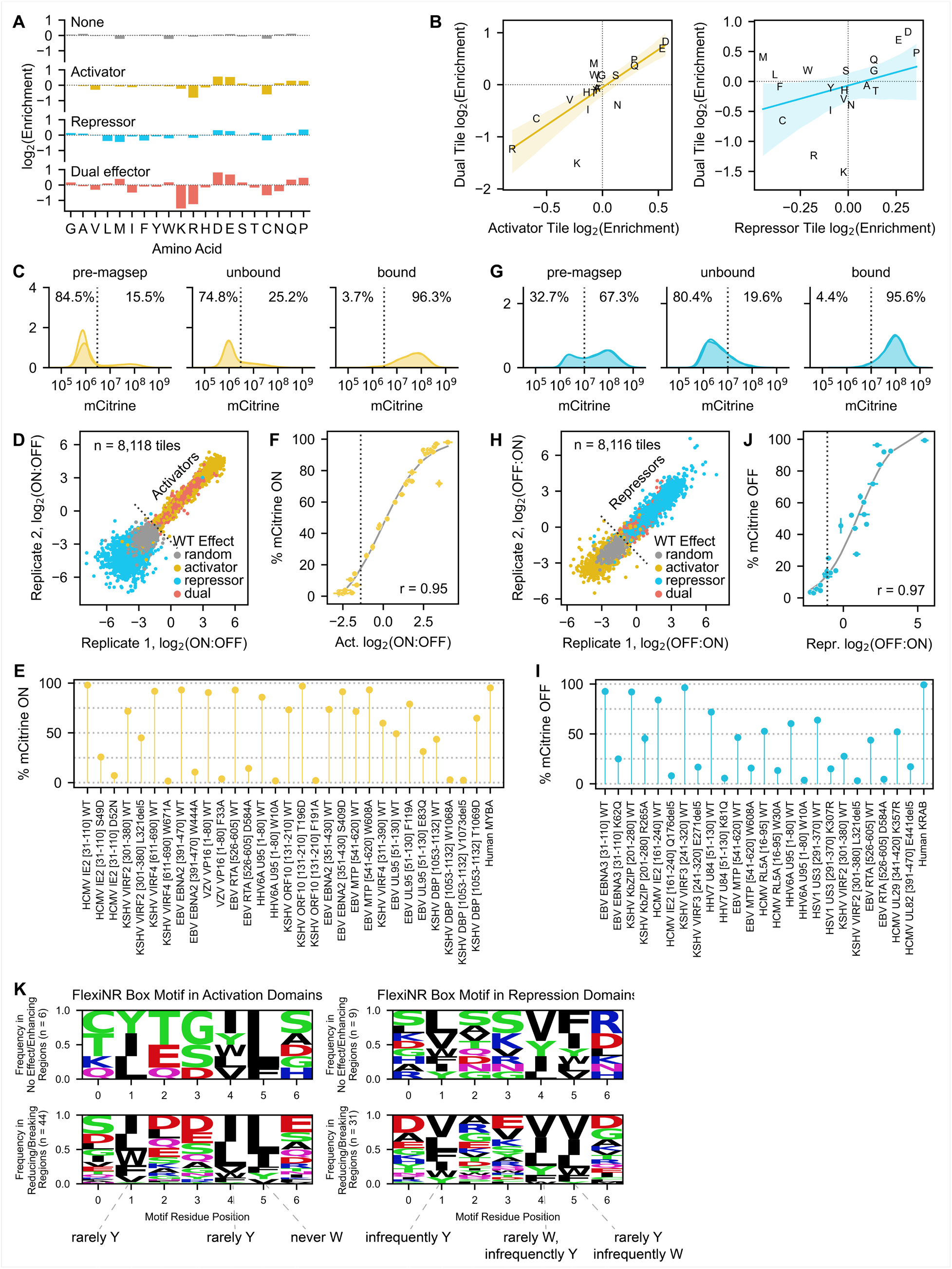
Herpesvirus sequence features and perturbation screen details and validation. (**A**) Barplots of the log2-transformed ratios of amino acid frequencies in tiles with no activity (gray, top row), activation (yellow, second row), repression (blue, third row), and dual effector (red, bottom row) tiles relative to their proteome frequencies. Positive values represent an enrichment in effector domains while negative values represent a depletion. (**B**) Comparison of the amino acid enrichments in (A) showing a stronger correlation between activator and dual effector tiles (left) than repressor and dual effector tiles (right). (**C**) Flow cytometry distributions on day two of recruitment for replicate screen minCMV reporter cells before and after magnetic separation. (**D**) Reproducibility of activation log2(ON:OFF) enrichment scores across two replicates, with each tile colored according to its original effector category in the HHV tiling screen (random, activator, repressor, and dual effector). (**E**) Summary of activation strengths of individually validated tiles as assessed by flow cytometry. Tiles are ranked by wild-type screen score, with percent of mCitrine-positive (ON) cells shown on the y-axis and the MFI of ON cells represented as marker size. (**F**) Relationship between activation log2(ON:OFF) enrichment score from the screen and the percent of ON cells for the set of individually validated tiles in (E). Logistic fit in gray with Spearman r = 0.95. The gray dashed line at x = -1.39 represents the detection threshold. (**G**) Flow cytometry distributions on day five of recruitment for replicate screen pEF reporter cells before and after magnetic separation. (**H**) Reproducibility of repression log2(OFF:ON) enrichment scores across two replicates. (**I**) Summary of repression strengths of individually validated tiles as assessed by flow cytometry. Tiles are ranked by wild-type screen score, with percent of mCitrine-negative (OFF) cells shown on the y-axis and the MFI of OFF cells represented as marker size. (**J**) Relationship between repression log2(OFF:ON) enrichment score from the screen and the percent of OFF cells for the set of individually validated tiles in (H). Logistic fit in gray with Spearman r = 0.97. The gray dashed line at x = -1.07 represents the detection threshold. (**K**) Logos showing the frequency of amino acids at each position of the flexiNR box motif for instances in regions whose deletion has no effect or enhances activity (top row) versus regions whose deletion reduces or breaks activity (bottom row) in activation (left) or repression (right) domains. The flexiNR motif used in this initial search was maximally flexible and was refined for subsequent analysis based on which amino acids rarely occurred at a given position (Methods). Negative charge and phosphorylatable residues (S/T) are common at the variable positions throughout this motif.

**Fig. S6.**
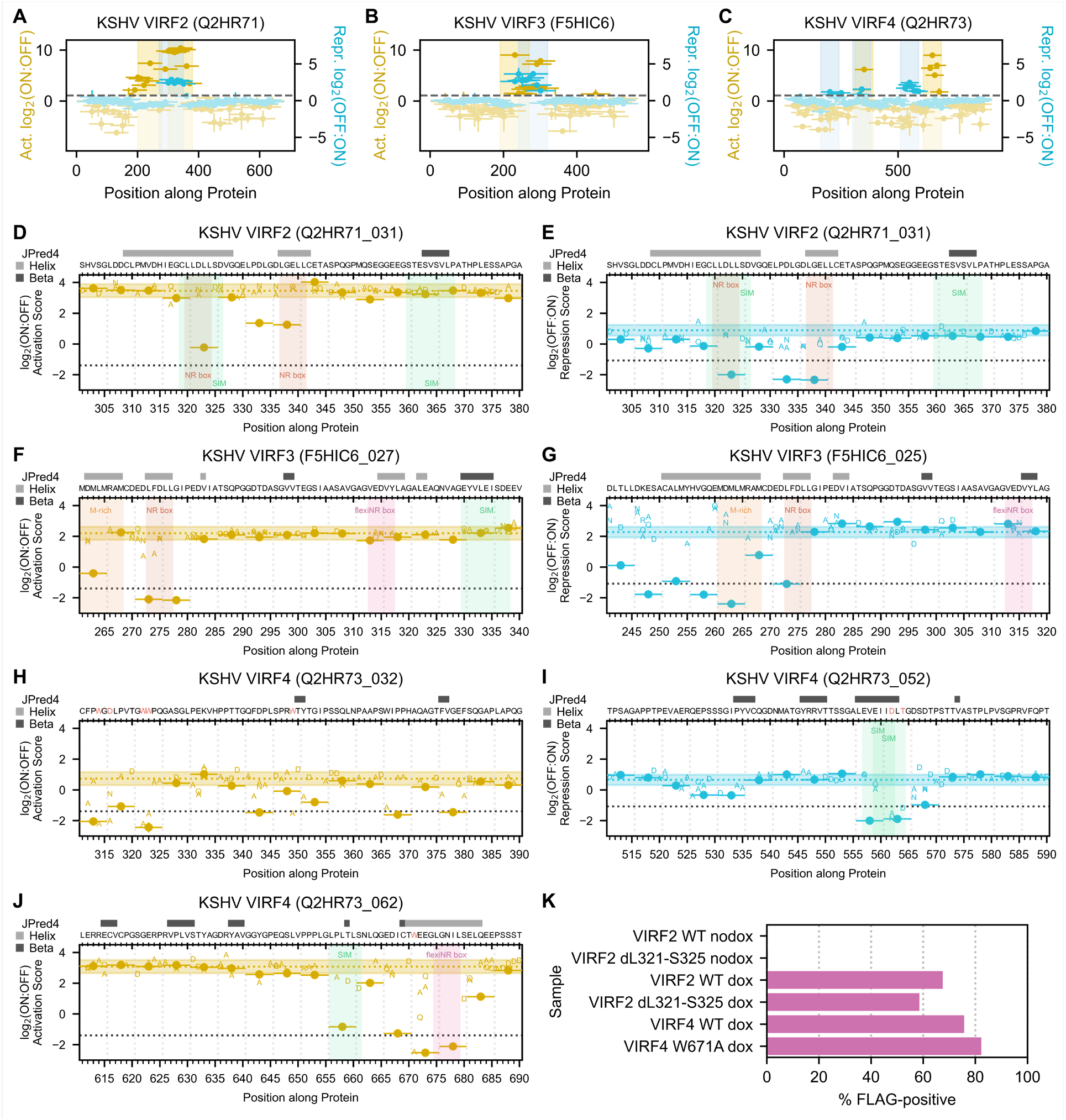
Identification of effector domains and essential amino acids in VIRF proteins. **(A-C)** Tiling plots of VIRF2 (A), VIRF3 (B), and VIRF4 (C), highlighting their activation (yellow), repression (blue), and dual (overlap of yellow and blue) effector domains. (**D-E**) Perturbation plots of the dual effector domain in VIRF2, showing the effects of 5aa deletions and single-residue substitutions on activation (D) and repression (E). Motifs are highlighted with a colored vertical span and indicated on the plot. (**F-G**) Perturbation plots of the dual effector domain in VIRF3, showing the effects on activation (F) and repression (G). (**H-J**) Perturbation plots of the dual effector domain (H, only activation shown), moderate-strength repression domain (I), and strong activation domain (J) in VIRF4. (**K**) Summary of 3xFLAG-tagged WT and mutant VIRF protein levels as estimated using anti-FLAG staining and flow cytometry. A dox-inducible promoter (minCMV driven by rTetR-VP48) is used to induce expression of the 3xFLAG-tagged full-length proteins indicated on the y-axes (See Fig. 6F, and Methods). Mutants: VIRF2 with deletion of L321 to S325 (dL321-S325) and VIRF4 with substitution of tryptophan 671 with alanine (W671A).

**Fig. S7.**
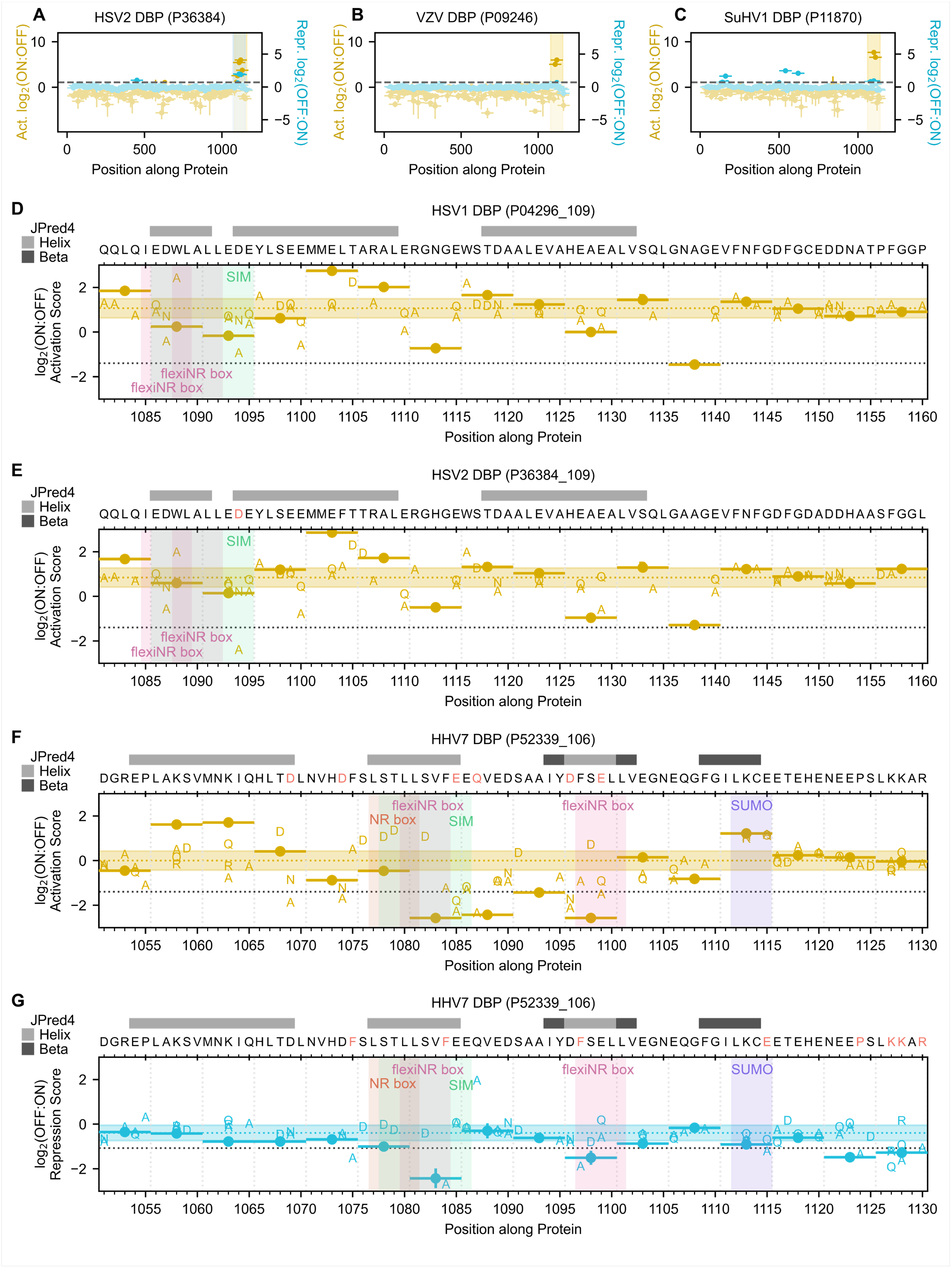
DBP domains and perturbation analysis. (**A-C**) Tiling plots of three alphaherpesvirus DBP homologs showing their C-terminal activation domains, which, in the case of HSV2 (A), also has repression potential. (**D-G**) Perturbation plots showing the effects of 5aa deletions and single-residue substitutions on activation (D, E, G) or repression (F) by various DBP homologs.

## Supplemental Tables

**Table S1.** Sequences and activation and repression screen measurements for the vTR-CoV tiling and Spike perturbation libraries compiled in an Excel file.

**Table S2.** Sequences and activation and repression screen measurements for the HHV tiling library compiled in an Excel file.

**Table S3.** Sequences and activation and repression screen measurements for effector domains identified in the vTR-CoV tiling and HHV tiling screens compiled in an Excel file.

**Table S4.** Sequences and activation and repression screen measurements for HHV perturbation and chemical inhibition screens (DMSO, SGC-CBP30 for p300/CBP inhibition, tazemetostat for EZH2 inhibition, and TMP269 for class IIa HDAC inhibition) compiled in an Excel file.

**Table S5.** RNA-seq Deseq2 analyses for the full-length viral proteins expressed in this study (each compared to a negative mCitrine expression control) compiled in an Excel File.

**Table S6.** Results of the initial and final motif search within herpesvirus effector domains compiled in an Excel File.

## Methods

### Plasmids used in this study

| Plasmid Code | Plasmid Description | Plasmid Name | Origin |
| --- | --- | --- | --- |
| pJT039 | Reporter to measure transcriptional repression | pJT039 AAVS1-PuroR-9xTetO-pEF-IGKleader-hlgG1_FC-Myc-PDGFRb-T2A-Citrine-PolyA | Tycko, et al. 2020. |
| pDY32 | Reporter to measure transcriptional activation | pDY32 AAVS1-PuroR-9xTetO-minCMV-IGKleader-hlgG1_FC-Myc-PDGFRb-T2A-Citrine-PolyA | Tycko, et al. 2020. |
| pJT126 | Lentiviral vector for recruiter expression | pJT126 lenti pEF-rTetR(SE-G72P)-3XFLAG-LibCloneSite-T2A-mCherry-BSD-WPRE | Tycko, et al. 2020. |
| pCL040 | Lentiviral vector for inducible transgene expression | pCL040 lenti TRE-3xFLAG-LibCloneSite pEF-rTetR(SE-G72P)-VP48-T2A-mCherry-BSD-WPRE | This study |

### Cell lines and cell culture

All experiments were performed with K562 cells (ATCC, CCL-243), which were cultured in RPMI 1640 (Gibco, 11-875-119) supplemented with 10% FBS (Omega Scientific, FB-15) and 1% Penicillin-Streptomycin-Glutamine (Gibco,10378-016). K562 reporter cell lines were the same as those used in the original HT-recruit study^8^ and were generated by TALEN-mediated homology-directed repair to integrate donor constructs (JT039 with EF1a reporter: Addgene #161927; DY032 with minCMV reporter: Addgene #161928) into the *AAVS1* locus using hAAVS1 1L TALEN (Addgene #35431) and hAAVS1 1R TALEN (Addgene #35432). These cell lines were not authenticated. HEK293T Lenti-X cells (Takara, 632180), which were used to package lentivirus as described in the following section, were cultured in DMEM with GlutaMAX (Gibco, 10-566-024) supplemented with 10% FBS and 1% Penicillin-Streptomycin (Gibco, 15-140-122). All cells were cultured in a controlled humidified incubator at 37°C and 5% CO_2_. All cell lines tested negative for mycoplasma.

### Lentiviral transduction

For small-scale lentivirus production, HEK293T Lenti-X cells were seeded into 6-well plates at 1 x 10^6^ cells per well in 2mL. The next day, cells were transfected with 750ng of an equimolar mixture of the three third-generation envelope and packaging plasmids (pMD2.G: Addgene #12259; pRSV-Rev: Addgene #12253; pMDLg/pRRE: Addgene #12251, all gifts from Didier Trono) and 750ng of lentiviral transfer plasmid encoding the transgene of interest after a 15 minute incubation of these plasmids with 10μL of polyethylenimine (PEI, Polysciences #23966). Lentivirus-containing culture supernatant was harvested 72 hours after transfection, passed through a 0.45μm PES filter (CELLTREAT #229749), and added undiluted to K562 cells for a final cell concentration of 3-4 x 10^5^ cells/mL for pJT126-based effector recruitment vectors (Addgene #161926) or 1-2 x 10^5^ cells/mL for pCL040-based inducible protein expression vectors (to be deposited on Addgene) to account for differences in infection efficiency. The cells were spinfected as follows: the cell-virus suspension was centrifuged in 15mL conicals at 1,000 x g for 2 hours at 33°C, after which the supernatant was removed from the cells (decanted), decontaminated, and discarded; cells were subsequently cultured for two days in fresh media to allow for integration and expression of mCherry and blasticidin resistance. Cells were treated with 10μg/mL Blasticidin S HCl (Gibco #A1113903) from days 2-10 post-infection to select for successfully transduced cells. Starting at day 3 post-infection, selection efficiency was monitored regularly by measuring mCherry positivity on a Bio-Rad ZE5 Cell Analyzer (12004278).

For screen-scale lentivirus production, HEK293T Lenti-X cells were seeded into 15-cm dishes at 13 x 10^6^ cells per dish in 30mL (approximately one dish per 5,000 library elements). The next day, cells were transfected with 11μg of an equimolar mixture of the three third-generation envelope and packaging plasmids and 11μg of the library of lentiviral transfer plasmids with 150μL of PEI. A full media change was performed 24 hours after transfection, and lentivirus-containing culture supernatant was harvested at 72 hours after transfection, applied to a 0.45μm PES filter unit (Thermo Scientific #1680045), and a fraction was used for titration to determine the appropriate dilution for approximately 25% mCherry-positive cells (equivalent to an MOI of 0.3 where approximately 90% of infected cells only receive one library member).

### vTR tiling library design

Protein sequences and metadata for the 419 human virus transcriptional regulators included in Table S2 of the vTR census study^7^ were downloaded from UniProt; however, only the 377 proteins from non-BSL4 viruses were considered for tiling due to safety concerns. Proteins tiles of 80aa in length were generated in 10aa increments along each protein, and duplicates were removed. Protein tile sequences were reverse-translated and codon-optimized using the Python package DNAchisel. Our codon optimization approach matched codon usage to natural human frequencies, excluded BsmBI sites, excluded C homopolymers greater than seven in length, enforced a local GC content of 20-70% within a 50bp window, and enforced an initial maximum global GC content of 65% that was incrementally relaxed by 1% if optimization failed. To the resulting 13,133 tiles, we added 1,500 80aa-long random negative controls whose codon usage matched natural human frequencies, and 386 additional sequences that would serve as fiducial controls across screens **(Table S1)**. To the 5’ and 3’ ends of all sequences, we appended BsmBI restriction sites for scarless Golden Gate cloning and library-specific primer handles for amplification by PCR, yielding a final length of 300nt for every oligonucleotide in the library. The vTR, CoV, and HHV libraries (design discussed below) were ordered as a single oligonucleotide pool from Twist Biosciences.

### CoV tiling library design

Protein sequences and metadata for the entire proteomes of 11 human and closely related bat coronaviruses were downloaded from UniProt, with most, but not all, of these sequences reviewed. For all ORF1a and ORF1ab polyprotein entries, we used the PTM/Processing > Chain information in UniProt to extract each of the individual non-structural protein sequences for tiling. For BtCoV-RaTG13 Orf1ab, which lacks Chain information in UniProt, we used the annotations of the near identical SARS-CoV-2 Orf1ab, accounting for the insertion of an isoleucine at residue 1023 for the SARS-CoV-2 homolog. Protein tiling, codon optimization, and appending restriction site and primer handle sequences were performed as above. The final CoV library comprised 7,564 unique coronavirus protein tiles, 850 80aa-long random negative controls, and 391 additional fiducial control sequences (**Table S1**).

### SARS-CoV-2 Spike perturbation library design

A multiple sequence alignment for all 11 full-length Spike homologs was performed with Clustal Omega to define the non-SARS-CoV-2 WT Spike sequences aligning to the repressive SARS-CoV-2 Spike-095 tile identified in the primary CoV tiling screen (UniProt P0DTC2, residues 941-1020). All other non-control library members were perturbations of the SARS-CoV-2 Spike-095 tile sequence and were generated as described in the main text. Altogether, the library comprised 9 WT protein sequences, 68 deletions, 69 consecutive double alanine substitutions, 64 consecutive triple alanine substitution, 760 single substitution elements in the ‘core’ region (residues 941-980), 100 non-consecutive double alanine substitutions within the ‘core’ region along the external trimer face, and one non-consecutive 15-residue alanine substitution of this same face. To assess the consequence of codon variation on protein function, three alternatively encoded oligonucleotides were designed for each of the unique WT sequence and perturbations described above, except for the SARS-CoV-2 WT Spike-095 sequence, which we alternatively encoded 10 different ways. The final library comprised 3,217 Spike-related elements, 381 80aa-long random negative controls from the vTR and CoV screens, and 100 additional fiducial control sequences. Codon optimization and appending restriction site and primer handle sequences were performed as above, except in the case of the deletion scanning elements, for which we added a filler sequence between the restriction site and primer handle sequence in order to maintain a uniform final oligonucleotide length of 300nt. This library (**Table S1**) was ordered as an oligonucleotide pool from Twist Bioscience.

### HHV tiling library design

Protein sequences and metadata for nearly the entire proteomes of 9 human herpesviruses were downloaded from UniRef90, which collapses UniProt entries on 90% sequence identity and represents each resulting protein cluster with a single, reviewed sequence, using the following search term: uniprot:(herpesvirus host:human NOT molluscum reviewed:yes) AND identity:0.9. A similar search on UniRef90 was performed for the Suid herpesvirus, which primarily infects pigs but is a commonly used model for studying alpha-herpesvirus biology. Two human herpesvirus protein sequences contained at least one X (HHV6B Q1: UniProt Q9QJ11; HHV6B Q2: UniProt P0DOE1), which required manual correction based on other entries in the cluster and the literature. Protein tiling, codon optimization, and appending restriction site and primer handle sequences were performed as above. The final library comprised 11,856 unique alpha-herpesvirus protein tiles, 13,679 unique beta-herpesvirus protein tiles, 7,434 unique gamma-herpesvirus protein tiles, 3,650 80aa-long random negative controls, and 413 additional fiducial control sequences (**Table S2**).

### HHV perturbation library design

Transcriptional effector domains identified in the primary HHV tiling screen were represented by their strongest tile. Screen scores were converted into estimated percent activation or repression based on the fit to the individual validation data described by the following logistic functions:

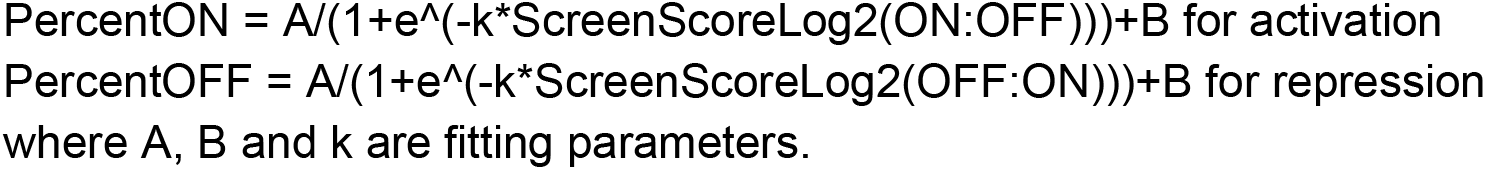

Only tiles whose percent activation or repression was estimated to be at least 40% (based on their screen scores and the equations above) were considered for perturbation in order to be able to measure appreciable differences in activity and to be able to test a larger set of perturbations for each tile. This criterion yielded 43 activator and 55 repressor tiles. While 21 of these activator and repressor tiles had some degree of dual effector activity, only eight met the 40% threshold for both activation and repression and were considered strong dual effector tiles. The protein-level deletions and substitutions described in the main text were generated with Python scripts, these sequences were reverse-translated and codon-optimized using DNAchisel, and restriction site and primer handle sequences were appended as above. Altogether, the library comprised 98 WT sequences, 1,567 deletions, and 6,129 substitutions: 268 F:A, 112 W:A, 171 Y:A, 557 D:A, 557 D:N, 621 E:A, 621 E:Q, 384 R:A, 218 K:A, 218 K:Q, 218 K:R, 645 S:D, 506 T:D, 327 Q:A, and 706 P:A. Given the low variability observed in the SARS-CoV-2 Spike perturbation screen between alternatively coded library members, each HHV perturbation library member was only encoded one way. To these 7,794 elements, we added 320 80aa-long random negative controls from the HHV tiling screens and 100 additional fiducial control sequences **(Table S4)**. This library was ordered as an oligonucleotide pool from Twist Bioscience.

### HHV hits library design for chemical inhibition screens

All library members that were above the activation or repression detection thresholds in the primary HHV tiling screens were included in the HHV hits library without modification (i.e. identical DNA sequence) except for the primer handle sequences. To the 194 activation hits, 74 dual effector hits, and 553 repression hits, we added 50 80aa-long random negative controls from the vTR and CoV screens and 71 additional fiducial control sequences. This library **(Table S4)** was ordered as an oligonucleotide pool from Twist Bioscience.

### Tiling and perturbation library cloning

Twist oligonucleotide pools were resuspended to a concentration of 10ng/μL in 10mM Tris-HCl pH 8.0 with 1mM EDTA. Individual libraries (e.g. vTR, CoV, HHV, etc.) were selectively PCR amplified using library-specific primers annealing to the primer handle sequences flanking each oligonucleotide. Between two to six 50μL PCR reactions were performed for each library to produce enough product for downstream cloning steps, and all reactions were prepared in a pre-PCR hood to mitigate DNA contamination. Each 50μL reaction consisted of 10μL of 5X Herculase II Reaction Buffer (Agilent #600675), 34.5μL of nuclease-free water, 0.5μL of 10ng/μL template (5ng total), 1μL of each 10μM primer, 1μL of DMSO, 1μL of 10nM dNTPs, and 1μL of Herculase II Fusion DNA Polymerase, added in that order. The thermocycling protocol was as follows: an initial denaturation at 98°C for 3 minutes; between 17 and 21 cycles of 98°C for 20s, 61°C for 20s, and 72°C for 30s; and a final extension at 72°C for 3 minutes. Initial small-scale test PCRs were performed to determine library-specific cycle numbers that yielded clean, visible amplicons suitable for gel extraction and not at saturation. Amplified oligonucleotide libraries were run on a 2% TBE gel, the 300bp bands were excised, and DNA was extracted from the agarose using the QIAquick Gel Extraction Kit (Qiagen #28704).

The pJT126 lentiviral recruitment vector (Addgene #161926) was pre-digested with 10,000 U/mL Esp3I (NEB #R0734L) at a ratio of 1μL of enzyme per 5μg of plasmid at 37°C for 15 minutes, followed by heat inactivation at 65°C for 20 minutes. Pre-digested pJT126 was run on a 0.5% TAE gel long enough to cleanly excise the digested product, which was subsequently extracted from the agarose. Oligonucleotide libraries were cloned into this vector using the GoldenGate cloning method, with between 10 to 16 20μL reactions per library. Each 20μL GoldenGate reaction consisted of 2μL of 10X T4 DNA Ligase Reaction Buffer (NEB #B0202S), nuclease-free water, 75ng of pre-digested pJT126, 5ng of amplified oligonucleotide library, and 2μL of NEBridge Golden Gate Assembly Kit (BsmBI-v2) (NEB #E1602L), added in that order. The NEBridge kit contains both BsmBI-v2 (an isoschizomer of Esp3I) and T4 DNA ligase. GoldenGate reaction conditions were 65 cycles of 42°C for 5 minutes then 16°C for 5 minutes, followed by a final digest at 42°C for 5 minutes and heat inactivation at 70°C for 20 minutes. Reactions were pooled, purified and concentrated with the MinElute PCR Purification Kit (Qiagen #28004), and eluted in 6μL of nuclease-free water.

Endura Electrocompetent Cells (Lucigen #60242-2) were thawed on ice for 10 minutes, then 25μL of cells were mixed with 2μL of the purified/concentrated GoldenGate product and transferred to a Gene Pulser/MicroPulser Electroporation Cuvettes, 0.1cm gap (Bio-Rad #1652089). Cells were electroporated on a Gene Pulser Xcell Total System (Bio-Rad #1652660) with the following conditions: 1.8kV, 10μF, 600Ω, and 0.1cm distance. Immediately after, 2mL of 37°C SOC Recovery Medium (NEB #B9020) were added to the cuvette, the contents of which were mixed by gentle pipetting and subsequently transferred to a 14mL round-bottom tube for a 1-hour recovery in a 37°C bacterial shaker. After recovery, cells were plated across four 10” x 10” luria broth agar plates with 100μg/mL carbenicillin, with a small amount of the recovery reserved for 1:100 dilution plating in triplicate to estimate library coverage. Plates were incubated in a warm room (approximately 33°C) for 14 to 18 hours, after which colonies were harvested by addition of luria broth and scraping. Cells were pelleted at 4,000 x g for 20 minutes, and plasmid pools were extracted using the Qiagen Plasmid Maxi Kit (Qiagen #12162). To assess library quality and representation bias, library members were amplified from the plasmid pool by PCR with primers containing Illumina adapters for readout by next generation sequencing.

### High-throughput recruitment assay with vTR, CoV, Spike perturbation, and HHV libraries

K562 reporter cells were infected with lentiviral libraries by centrifugation at 1,000 x g for 2 hours. Infection was performed with two replicates per library and the following number of starting cells per replicate per reporter line: 45 x 10^6^ cells for the pooled vTR and CoV libraries; 12.5 x 10^6^ cells for the Spike perturbation library; 45 x 10^6^ cells for the HHV tiling library; 15 x 10^6^ cells for the HHV perturbation library; and 2.5 x 10^6^ cells for the HHV hits library screened in the presence of chemical inhibitors. Estimates of infection coverage (the average number of cells infected with a given library member) are as follows: 420X and 330X for the minCMV and EF1a reporter lines, respectively, for the pooled vTR and CoV libraries; 900X for the EF1a reporter line infected with the Spike perturbation library; 320X and 290X for the minCMV and EF1a reporter lines, respectively, infected with the HHV tiling library; 250X and 200X for the minCMV and EF1a reporter lines, respectively, infected with the HHV perturbation library; and 250X and 350X for the minCMV and EF1a reporter lines, respectively, infected with the HHV hits library. Cells were treated with 10μg/mL blasticidin (Gibco #A1113903) starting two days after infection for approximately five to seven days total when at least 80% of cells were mCherry positive as monitored by daily flow cytometry (cells were analyzed no earlier than three days after infection in compliance with safe lentivirus practice).

For the vTR/CoV and HHV libraries, cells were maintained in 1L spinner flasks with constant, gentle paddle rotation. For the Spike perturbation, HHV perturbation, and HHV hits libraries, cells were maintained in vented T175, T225, and T25 flasks, respectively. All cultures were maintained in log growth conditions with daily half-volume media changes to dilute cells back to approximately 5 x 10^5^ cells/mL, making sure to never drop the maintenance coverage (the number of cells harboring a given library member) below the initial infection coverage. Following antibiotic selection, recruitment was induced by treating the cells with 1000ng/mL doxycycline hyclate (Tocris #4090) for two days for activation screens or for five days for repression screens. Half the amount of doxycycline was replenished each day under the assumption of a 24-hour half-life. For screens with the HHV hits library, cells were treated with 10μM SGC-CBP30 (Selleck Chemicals #S7256), 10μM tazemetostat (Selleck Chemicals #S7128), 10μM TMP269 (Selleck Chemicals #S7324), or DMSO (vehicle) for 24 hours prior to dox addition and throughout the recruitment timecourse, with these chemical inhibitors replenished daily during media changes.

### Magnetic separation for high-throughput recruitment assays

At assay endpoint, a volume of cells equivalent to 10,000X coverage was pelleted and washed twice with DPBS (Gibco #14190-250) to remove immunoglobulins from the FBS. Cell pellets were resuspended in magnetic separation wash buffer (2% BSA in DPBS) at a concentration of 23 x 10^6^ cells/mL, and a small volume was reserved as an ‘input’ sample for analysis by flow cytometry (described below). In parallel, a volume of paramagnetic Dynabeads M-280 Protein G (Thermo Fisher, #10003D) of 3-9μL per 1 x 10^6^ cells (scaled based on the rarity of the bound population) were diluted in five volumes of wash buffer, incubated on a magnetic stand for 2 minutes, cleared of the supernatant, and resuspended in the cell suspension. The cell-bead suspension was incubated at room temperature for 75 minutes on a nutator to allow adequate time for cells expressing the IgG surface marker to bind the protein G-functionalized Dynabeads, and the suspension was subsequently incubated on a magnetic stand for 5 minutes to separate bead-bound and unbound fractions. The unbound fraction was transferred to a new tube, which was incubated again on a magnetic stand for 5 minutes to clear the suspension of any remaining beads, and the unbound fraction was transferred to a final tube. All beads in the original and second tube were pooled by resuspending in a volume of wash buffer equivalent to the initial volume. This tube was incubated at room temperature for 15 additional minutes on a nutator, subsequently incubated on a magnetic stand for 5 minutes, and the unbound ‘wash’ fraction was transferred to a new tube. The remaining bead-bound cell fraction was resuspended in a volume of wash buffer equivalent to the initial volume, and all three fractions (unbound, wash, and bead-bound) as well as the input sample were run on a Bio-Rad ZE5 Cell Analyzer to assess the effectiveness of magnetic separation and to estimate the total number of cells recovered in each fraction. As expected with the dual surface marker-mCitrine reporter, the unbound fraction had low mCitrine and the bead-bound fraction had high mCitrine. In every screen, the wash fraction (typically less than 5% of the total sample) mCitrine distribution resembled that of the input sample, and thus this fraction was discarded. The unbound and bead-bound fractions were pelleted by centrifugation at 600 x g for 5 minutes, decanted, and frozen at -20°C.

### Library preparation and sequencing

For all high-throughput recruitment assays, genomic DNA was extracted for pelleted cell fractions with one of the following: DNeasy Blood & Tissue Kit (Qiagen #69504) for fractions with fewer than 5 x 10^6^ cells; QIAamp DNA Blood Midi Kit (Qiagen #51183) for fractions with 5-20 x 10^6^ cells; and Blood & Cell Culture DNA Maxi Kit (Qiagen #13362) for fractions with 20-100 x 10^6^ cells. During genomic DNA extraction, bead-bound fractions were incubated on a magnetic stand to remove beads prior to loading lysate onto the silica columns. Genomic DNA was eluted in Buffer EB (Qiagen #19086) rather than the provided Buffer AE (Qiagen #19077) to avoid inhibition of PCR.

Library members were amplified by PCR with primers containing Illumina adapter extensions at a final concentration of 500nM. All PCRs were prepared in a pre-PCR hood to reduce the likelihood of contamination by amplicons and plasmids. Small-volume test PCRs across a range of cycle numbers were performed and visualized by gel electrophoresis to identify the optimal cycle number that yielded sufficient material for extraction without reaching saturation and without producing non-specific bands. Final DNA template concentrations of 100-200ng/μL were used when possible and standardized for a given fraction (unbound or bound) across screen replicates, and at least one-third of all extracted genomic DNA was used as input for PCR to preserve library representation, resulting in a variable number of PCRs for each fraction. PCRs for screens with the vTR and CoV libraries were performed using NEBNext High-Fidelity 2X PCR Master Mix (NEB #M0541L) with 33 cycles. PCRs for screens with all other libraries were performed using the NEBNext Ultra II Q5 Master Mix (NEB #M0544L) with 21 to 24 cycles. Thermocycling and subsequent steps were performed outside of the pre-PCR hood. The thermocycling protocol was as follows: an initial denaturation at 98°C for 3 minutes; the aforementioned number of cycles of 98°C for 10s, 63°C for 30s, and 72°C for 30s; and a final extension at 72°C for 3 minutes. All reaction products for a given fraction were pooled and mixed, and 150μL were subsequently run on a 1% TAE gel, extracted, purified using the QIAquick Gel Extraction Kit, and eluted with 30μL of Buffer EB. The concentrations of each sample were quantified with the Qubit dsDNA HS Assay Kit (Thermo Fisher #Q32854) on a Qubit 4 Fluorometer (Thermo Fisher #Q33238), pooled with 15% PhiX Control v3 (Illumina #FC-110-3001), and sequenced one of the following ways: on an Illumina NextSeq 550 with 1 x 75 or 1 x 150 cycles, on an Illumina HiSeq 2000 with 2 x 150 cycles, or on an Illumina MiSeq with 2 x 150 cycles.

### Sequencing analysis

Sequencing data was processed and analyzed using the HT-recruit Analyze code first described in the original study^8^ and available on GitHub (https://github.com/bintulab/HT-recruit-Analyze). Briefly, reads were demultiplexed with bcl2fastq (Illumina), aligned with ‘makeCounts.py’ to a reference (made using ‘makeIndices.py’), and used to compute enrichment scores between unbound (OFF) and bound (ON) fractions for each library member using ‘makeRhos.py’. Library members with fewer than five reads in both fractions for a given replicate were filtered out, while those with fewer than five reads in one fraction had their reads adjusted to five reads for that fraction to avoid inflation of enrichment scores. For all screens, library members with a sum of fewer than 50 reads between both fractions for both replicates were filtered out, as these would produce noisy enrichment scores. For all screens, the detection threshold above which we estimated we could measure transcriptional effector activity was set at two standard deviations above the mean enrichment score of the negative random control population.

### Individual recruitment assay validations by flow cytometry

Library members selected for individual validation experiments were ordered as gene fragments from Twist or IDT and cloned into the pJT126 lentiviral recruitment vector using Golden Gate cloning. Lentivirus was prepared and used to transduce reporter cells in replicate as described above. Following selection, cells were split into two wells, one of which was untreated and the other treated with 1μg/mL doxycycline for 2 days for the activation assay or 5 days for the repression assay. Half-media changes were performed daily, replenishing doxycycline for the appropriate wells, and 10,000 cells from each well were passed through a 40μm filter to be analyzed on a Bio-Rad ZE5 Flow Analyzer daily to monitor changes to reporter transcriptional state. Data was analyzed using Cytoflow^61^, first gating events for viability and mCherry expression. Gates for mCitrine expression were set based on an rTetR-only negative control to compute the fraction of mCitrine ON and OFF cells on each day of doxycycline treatment. Additional analyses and visualizations were performed with custom Python scripts.

### Estimation of 3xFLAG-tagged protein levels by anti-FLAG staining and flow cytometry

All effector recruiter fusions (pJT126 vector) and full-length viral proteins for inducible expression (pCL040 vector) were designed as fusions to a 3xFLAG epitope tag to enable estimation of protein levels by anti-FLAG staining. Briefly, Fix Buffer I (BD Biosciences #557870) was pre-warmed to 37°C for 15 minutes, and Perm Buffer III (BD Biosciences #558050) was pre-chilled on ice. Approximately 1 x 10^6^ cells were harvested by centrifugation at 300 x g for 5 minutes and washed once with DPBS. Cells were resuspended in 50uL of Fix Buffer I, incubated at 37°C for 15 minutes for fixation, pelleted by centrifugation, and washed once with 500μL cold DBPS with 10% FBS. Cells were resuspended in 50uL of Perm Buffer III, incubated on ice for 30 minutes for permeabilization, pelleted by centrifugation, and washed once with 500μL cold DBPS with 10% FBS. Cells were resuspended in an antibody solution containing 5μL DYKDDDDK Epitope Tag Alexa Fluor 647-conjugated Antibody (R&D Systems #IC8529R) and 45μL DBPS + 10% FBS and incubated in the dark at room temperature for one hour. Cells were pelleted by centrifugation, washed with 500μL cold DBPS with 10% FBS, resuspended in 250μL DBPS with 10% FBS, and filtered through a 40μm filter prior to analysis on a Bio-Rad ZE5 Flow Analyzer. Data was analyzed using Cytoflow, first gating samples for viability, and, in the case of effector recruiter fusions, for mCherry expression. Gates for FLAG positivity were set based on wild-type or uninduced cells lacking the 3xFLAG epitope. Additional analyses and visualizations were performed with custom Python scripts/

### Multiple sequence alignment

Protein sequences to be aligned were compiled into a single FASTA file either manually or by querying multiple identifiers in UniProt and downloading a FASTA file of the compiled results. These files were run through Clustal Omega (https://www.ebi.ac.uk/Tools/msa/clustalo/) with the default settings and the ‘ClustalW with character counts’ output format. Alignment files were downloaded and either visualized in JalView or with custom python scripts using Biopython.

### Bulk RNA-seq

Gene fragments encoding full-length wild-type or mutant viral proteins were ordered from Twist or IDT and cloned into the pCL040 lentiviral inducible expression vector. Lentivirus was prepared and used to transduce wild-type cells in replicate as described above. Following selection, cells were cultured in 12-well plates and treated with 1μg/mL doxycycline to induce expression of viral transgenes. On day 2 post-induction, approximately 1 x 10^6^ cells were harvested by centrifugation at 300 x g for 5 minutes. RNA was extracted using the RNeasy Mini Kit (Qiagen #74104), with a volume of 600μL Buffer RLT for cell lysis and with the QIAshredder columns (Qiagen #79654) for lysate homogenization. For all samples, the RNA integrity number was 10 as assessed by the Stanford Protein and Nucleic Acid (PAN) Biotechnology Facility using the RNA Nano Kit (Agilent #5067-1511) on an Agilent Bioanalyzer. A total of 500ng of purified RNA was used as input for the NEBNext Ultra II RNA Library Prep Kit (NEB #E7770S), which first involved enrichment of polyadenylated mRNA using the NEBNext Poly(A) mRNA Magnetic Isolation Module (NEB #E7490). All steps were performed in accordance with the NEB protocol, with nine PCR cycles used for library amplification. Library size distributions were determined using the High Sensitivity DNA Kit (Agilent #5067-4626), and sample concentrations were quantified using the Qubit dsDNA HS Assay Kit on a Qubit 4 Fluorometer. Samples were pooled at equimolar ratios and sequenced on either a NextSeq 550 with 2 x 37 cycles or a MiSeq with 2 x 150 cycles.

Sequencing reads were demultiplexed with bcl2fastq. A FASTA of the GRCh38 human reference genome build was modified to include the viral transgenes of interest as separate chromosomes. The resulting FASTA was used to construct both a custom reference transcriptome using hisat2-build and a custom GTF genome annotation file using the script ‘make_transgene_gtf.py’. Paired reads were aligned to the custom reference using hisat2, and output SAM files were converted to BAM files using samtools. A differential expression analysis was performed in R with the Bioconductor DESeq2 package using a set of custom R scripts that were largely based on the workflow and commands described in the following tutorial: http://bioconductor.org/help/course-materials/2016/CSAMA/lab-3-rnaseq/rnaseq_gene_CSAMA2016.pdf. Additional analyses and data visualization were performed in python with custom scripts.

### Motif finding

Regular expressions describing short linear motifs (SLiMs) associated with gene regulation were pulled from the Eukaryotic Linear Motif (ELM) resource (http://elm.eu.org/) and used by custom Python scripts for pattern matching within protein sequences of interest. For the HHV perturbation screen data, an initial search with 40 motifs was conducted, specifically aimed at comparing motif frequencies in regions whose deletion either 1) had no effect or enhanced effector activity, or 2) reduced or completely broke effector activity. Sixteen motifs were found at a higher rate within the latter category and were used for a second search focused on annotating the overlap between these motifs and the effector domain essential regions (those whose deletion completely breaks activity) to identify potential cofactors **(Table S6)**.

The initial motif search included a new motif that we termed the flexiNR box based on its similarity to the traditional NR box motif (LxxLL). This motif was included on the basis of reported flexibility of the NR box in other human proteins^39–42^ and our own observations when examining the data, and it initially tolerated V, L, I, W, F, or Y at every position in the original NR box motif containing an L. The regular expression in the initial search was: ([^∧^P][VIWFY][^∧^P][^∧^P][VLIWFY][VLIWFY][^∧^P])|([^∧^P][VLIWFY][^∧^P][^∧^P][VIWFY][VLIWFY][^∧^P])|([^∧^P][VLIWFY][^∧^P][^∧^P][VLIWFY][VIWFY][^∧^P]).

Logos of the motif instances in the no effect/enhancing regions versus reducing/breaking regions in activation and repression domains were generated using the ‘logomaker’ Python package. From these, we determined that position 1 of the motif rarely contained Y, position 4 rarely contained W, and the position 4 rarely contained W or Y, resulting in the final pattern: ([^∧^P][VIWF][^∧^P][^∧^P][VLIFY][VLIF][^∧^P])|([^∧^P][VLIWF][^∧^P][^∧^P][VIFY][VLIF][^∧^P])|([^∧^P][VLIWF][^∧^P][^∧^P] [VLIFY][VIF][^∧^P]).

### CRISPR/Cas9 targeting of the KSHV DBP gene (ORF6)

CRISPR/Cas9 sgRNAs were introduced to cells and their effects on late gene expression and replication were measured as previously described in^51^. Briefly, HEK293T and iSLK cells were grown in DMEM (Gibco, +glut, +glucose, -pyruvate) with 10% FBS (Peak Serum), pen-strep (Gibco), and additional 1X GlutaMAX (Gibco). The Cas9+ iSLK line latently infected with a version of the BAC16 KSHV genome^62^ modified to contain a reporter of late gene activity (K8.1 promoter driving an mIFP2 fluorescent cassette) was maintained in 1 μg/mL puromycin, 50 μg/mL G418, 10 μg/mL blasticidin, and 125 μg/mL hygromycin.

sgRNAs targeting along the ORF6 gene were cloned into a mU6-driven guide expression plasmid (Addgene #89359) and delivered via lentiviral transduction at high MOI to the Cas9+ iSLK-BAC16-K8.1pr-mIFP2 cells. To analyze late gene expression, cells were were treated with 5 μg/mL doxycycline, which induces expression of KSHV RTA (ORF50) for lytic reactivation, and 1 mM sodium butyrate, an HDAC inhibitor that facilitates reactivation. Forty-eight hours after reactivation, cells were fixed in 4% PFA and quantified using flow cytometry (BD LSRFortessa). To analyze viral DNA replication, cells were similarly reactivated, and 48 hours afterwards were treated with 30 μM EdU for two hours. Cells were then trypsinized and fixed using 4% PFA. EdU was labeled with Cy5 using the Click-IT flow cytometry kit (Invitrogen) and quantified by flow cytometry (BD Accuri C6 Plus). All experiments were performed in four replicates from different days and independent reactions.

To monitor ORF6 mRNA levels, cells were reactivated, and 24 hours post-reactivation, RNA was harvested using RNA QuickExtract (Lucigen). Turbo DNAse (Invitrogen) was used to remove DNA. AMV RT (Promega) with 9 bp random primers was used for reverse transcription. qPCR was performed using iTAq Universal SYBR Green (Bio-Rad) amplifying the coding region of ORF6 along with primers targeting the host 18S RNA.

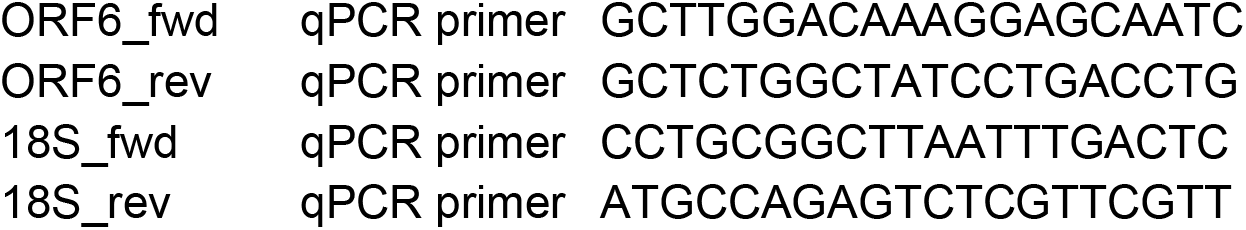

To monitor editing at the ORF6 locus, gDNA was extracted from unreactivated cells using DNA QuickExtract. The ORF6 locus was amplified using GoTaq (Promega) and Sanger sequenced. Editing outcomes were then determined with the Synthego ICE software.

## Acknowledgements

We thank Michaela M. Hinks and Benjamin R. Doughty for sequencing assistance, Adi Mukund for data visualization advice, and members of the Bintu lab for helpful conversations and assistance. We thank Twist Biosciences for oligonucleotide library synthesis. This work was supported by NIH-NIGMS MIRA R35GM12894701 (LB), NIH-NHGRI R01HG011866 (LB, MCB), NIH T32GM145402 (ART), NIH-NIAID R01AI122528 (BAG), Sarafan Chem-H Chemistry-Biology Interface Training Grant (ART), Howard Hughes Medical Institute and Life Sciences Research Foundation Postdoctoral Fellowship (DWM), and NIH-4K00DK126120-03 (JT). BAG is an investigator of the Howard Hughes Medical Institute.

## Author Contributions

CHL and LB designed the study, with significant intellectual contributions from ART. CHL designed the libraries. CHL and ART cloned the libraries. CHL performed the vTR/CoV tiling, HHV tiling, HHV perturbation, and HHV chemical inhibition screens. ART performed the Spike perturbation screen. CHL analyzed all screen data. CHL and ART performed RNA-seq and analyzed the data. CHL and ART generated plasmids and cell lines and performed individual recruitment assay experiments. DWM and KJY performed CRISPR/Cas9 experiments with KSHV and analyzed the data. CHL analyzed all other data, with contributions from ART and LB. CHL and LB wrote the manuscript, with significant contributions from ART and contributions from DWM and BAG. JT and MCB provided technical advice and materials at the beginning of the project. LB supervised the project.

